# Transcriptomic analysis of seed development in *Paysonia auriculata* (Brassicaceae) identifies genes involved in hydroxy fatty acid biosynthesis and seed maturation

**DOI:** 10.1101/2022.09.06.506826

**Authors:** Hao Hu, Armond Swift, Margarita Mauro-Herrera, James Borrone, Guadalupe Borja, Andrew N. Doust

## Abstract

*Paysonia auriculata* (Brassicaceae) produces multiple hydroxy fatty acids as major components of the seed oil. We tracked the changes in seed oil composition and gene expression during development, starting 14 days after flowers had been pollinated. Seed oil changes showed initially higher levels of saturated and unsaturated fatty acids (FAs) but little accumulation of hydroxy fatty acids (HFAs). Starting 21 days after pollination (DAP) HFA content sharply increased, and reached almost 30% at 28 DAP. Total seed oil also increased from a low of approximately 2% at 14 DAP to a high of approximately 20% by 42 DAP. We identified almost all of the fatty acid synthesis and modification genes that are known from Arabidopsis, and, in addition, a strong candidate for the hydroxylase gene that mediates the hydroxylation of fatty acids to produce valuable hydroxy fatty acids (HFAs) in this species. The gene expression network revealed is very similar to that of the emerging oil crop, *Physaria fendleri*, in the sister genus to *Paysonia*. Phylogenetic analyses indicate the hydroxylase enzyme, FAH12, evolved only once in *Paysonia* and *Physaria*, and that the enzyme is closely related to FAD2 enzymes. Phylogenetic analyses of FAD2 and FAH12 in the Brassicaceae and outgroup genera suggest that the branch leading to the hydroxylase clade of *Paysonia* and *Physaria* is under relaxed selection, compared with the strong purifying selection found across the FAD2 lineages.

## 1 Introduction

*Paysonia auriculata* (Engelm. & A. Gray) O’Kane & Al-Shehbaz is a member of a small genus of mustards that, along with its sister genus *Physaria*, contain hydroxy fatty acids in their seed oils. These two genera were formed from the reorganization of the genera *Lesquerella* and *Physaria* (Al-Shehbaz and O’Kane, 2002) and have recently been included in a larger phylogeny of the tribe Physareae, where their monophyly and relationships have been confirmed (Fuentes-Soriano and Kellogg, 2021). Hydroxy fatty acids (HFAs) are not known to occur elsewhere in the Brassicaceae, and are found only sporadically throughout the flowering plants, suggesting they have been derived multiple times from more common precursors (Napier, 2007). Because of the -OH group and the extra double bonds, HFAs are high-value oleochemicals with broad industrial applications, such as surfactants, lubricants, cosmetics, and antibacterial agents, and are the precursors for the synthesis of biodegradable bioplastics, polyhydroxyalkanoates, and additives for enhancing lubricity of biodiesel (Carlsson et al., 2011;Beaudoin et al., 2014;Aznar-Moreno and Durrett, 2017). However, much of the HFAs used in industry are sourced from oil from the seed of the castor bean plant (*Ricinus communis*), which contains highly toxic compounds (ricin and agglutinin) and strong allergens (2S albumin and 11S globulin) (Lago, 2009;Ashraf Ashfaq et al., 2018). Because of this, the production of castor oil is banned in the United States, and the U.S. imports many thousands of tons of castor oil from developing countries such as India (over 60,789 tons of castor oil in 2017, worth $96 million U.S. dollars (FAOSTAT, Food and Agriculture Organization Corporate Statistical Database)). Due to a rapid increase in demand, the global castor oil market size is anticipated to reach over $2 billion U.S. dollars by 2025 (www.marketwatch.com). Both social and scientific imperatives have spurred the search for safer alternatives to castor oil production, including identification and development of HFA production in other species. Both *Physaria* and *Paysonia* species produce a variety of HFAs, of which some, such as lesquerolic acid in *Physaria fendleri*, have receive significant attention, whereas others, such as those found in *Paysonia* species, have not. In addition, there has been little study in *Paysonia* of the genetic pathways underlying HFA production.

Castor oil contains very high levels (90%) of ricinoleic acid (18:1-OH) (Mutlu and Meier, 2010), while seeds of *Physaria* and *Paysonia* produce lower quantities (33-66%) of several different HFAs (Salywon et al., 2005). *Physaria* species and two species of *Paysonia* (*P. grandiflora* and *P. lasiocarpa*) produce lesquerolic acid (20:1-OH), the Tennessee/Alabama clade of *P. perforata, P. lyrata, P. densipila, P. lescurii*, and *P. stonensis* produce predominantly densipolic acid (18:2-OH) (Salywon et al., 2005), and *P. auriculata* produces the most energy rich, auricolic acid (20:2-OH), a HFA with one hydroxyl group and two double bonds. The percentage of oil that is hydroxylated in both *Physaria* and *Paysonia* species is less than that in castor oil presumably because not all positions in the triacylglyceride (TAG) in *Paysonia* and *Physaria* are occupied by hydroxy fatty acids (Cruz and Dierig, 2015). HFAs of both *Physaria* and *Paysonia* have an o-6-hydroxy-o-9-ene structure similar to ricinoleic acid from castor oil, and the seeds are biologically toxin free, so can be considered to be safe alternative sources of HFAs (Reed et al., 1997;Chen et al., 2011).

The biosynthesis of HFAs have been extensively studied in castor, *Physaria*, and Arabidopsis (Lee et al., 2015), and several lines of evidence suggest that the hydroxylation of FAs to become HFAs occurs mainly within the endoplasmic reticulum (ER), via the complicated cross-talks and interconversions between triacylglycerol (TAG) incorporation (Kennedy pathway), and phosphatidylcholine (PC) remodeling (acyl editing and Lands cycle) (Bates and Browse, 2012;Chapman and Ohlrogge, 2012;Vanhercke et al., 2013). Both castor and *P. fendleri* share the same route for hydroxylating 18:1-PC into 18:1-OH-PC, using Δ12 oleic acid hydroxylase (*FAH12*) to catalyze hydroxylation before being exported to the cytosol from ER as 18:1-OH, and then being activated to 18:1-OH-CoA. In castor, the 18:1-OH-CoA rejoins the Kennedy pathway, and finally forms 18:1-OH TAG (Bafor et al., 1991). In *P. fendleri*, 18:1-OH-CoA is further elongated into 20:1-OH-CoA by 3-ketoacyl-CoA synthase 18 (*PfKCS18*), and ultimately produces 20:1-OH TAG (Moon et al., 2001). *Physaria fendleri* also accumulates a small amount of other HFAs. In addition to *FAH12*, Δ12 oleic acid desaturase (*FAD2*) and/or Δ15 linoleic acid desaturase are hypothesized to be involved in the modification of FAs to produce HFAs (Engeseth and Stymne, 1996;Reed et al., 1997;Smith et al., 2003). In contrast, there has been little investigation of the genetic control of production of the different HFAs in *Paysonia*, although the study by Reed et al. (1997) suggested that the same enzymes (desaturases, elongases, hydroxylases) are present with possible modifications in the order of reactions.

The function of the hydroxylase in castor and in *Physaria* is similar in that they are both responsible for Δ12 hydroxylation to produce 18:1-OH (van de Loo et al., 1995). However, *P. fendleri PfFAH12* differs from castor *RcFAH12* by being bifunctional, catalyzing both Δ12 hydroxylation to produce 18:1-OH as well as Δ12 desaturation to produce 18:2-OH (Broun et al., 1998a). The bifunctional nature of the *PfFAH12* and the close sequence similarity, active diiron catalysis site and alignment of histidine residues between the hydroxylase and desaturase in both *P. fendleri* and castor further suggests the close relationship between these two enzyme classes (Broun et al., 1998a;Shanklin and Cahoon, 1998).

Analysis of the timing of HFA production in castor and *Physaria* seed development has shown that most HFA accumulation occurs after embryogenesis but before the maturation and eventual desiccation of the seed (Chen et al., 2009;Chen et al., 2011). This middle stage of seed development, where the embryo has been formed and is actively growing, is characterized in *Arabidopsis* by the change in color from the pale yellow color of the seed undergoing embryogenesis to green as photosynthetic pigments accumulate (Baud and Lepiniec, 2009). In both *P. fendleri* and *Physaria lindheimeri*, this stage occurs between 28-35 days after pollination (DAP), and is accompanied by rapid synthesis and accumulation of TAG and hydroxy fatty acids (Chen et al., 2011;Chen et al., 2017). Transcriptome analysis of *P. fendleri* around 30 DAP showed high expression of genes involved in fatty acid synthesis and modification, and high levels of conservation of those genes, apart from the hydroxylase, between *Physaria* and Arabidopsis, based on the annotations from AraLip and AraCyc databases (Horn et al., 2016).

In contrast to *Physaria*, there has been no study of the genes involved in HFA production in *Paysonia* species, although a study of the timeline of seed development and HFA oil production by Reed et al. (1997) suggests that HFA production starts as early as 14 DAP in *Paysonia* (the *Paysonia* ‘species’ assayed under its former genus name, *Lesquerella*, was *L. ‘kathryn’*, a hybrid of several Tennessee *Paysonia* species). This is much earlier than in *Physaria*, where HFA production was shown to commence approximately 40 DAP. We chose to develop a transcriptional developmental timeline for the Oklahoma species of *Paysonia, P. auriculata*, because it has the most complex HFA that is accumulated in large quantities, with two double bonds in addition to the hydroxyl group, and because *P. auriculata* appears to be the first species within the genus to possess a predominantly different HFA from both *Physaria* and the two most basal members of *Paysonia, P. grandiflora* and *P. lasiocarpa* (Salywon et al., 2005;Borja, 2013;Fuentes-Soriano and Kellogg, 2021). We constructed a developmental timeline for seed maturation and sampled weekly after pollination in order to explore transcriptome profile changes and HFA composition changes over the maturation period of the seed. We also used the transcriptome data to investigate the evolution of several key enzymes in the fatty acid modification pathway.

## 2 Materials and Methods

### 2.1 Plant materials, crossing and sample collecting

*P. auriculata* seeds were collected from a population of the species on the shores of the Salt Plains Reservoir, Alfalfa County, Oklahoma (Borja 1021, voucher in Oklahoma State University Herbarium). Seeds were germinated in BM2 Berger Plug Soil (Berger, Quebec City, Canada), transferred as seedlings to 3.8L pots filled with Sungro 380 Soil (Sungro, MA), and grown in a walk-in controlled growth room at temperatures between 28°C (day) and 22°C (night), with 16 hours lighting. After four weeks, young plants at the 8-12 leaf stage were transferred to a greenhouse with temperatures between 24° and 30°C, with daylight supplemented by metal halide lighting, for a photoperiod of 15 hours. Fertilizer (Jack’s 20:10:20, NPK, J.R. Peters, PA) was applied as an aqueous solution weekly.

Flowering and crosses between accessions started approximately four weeks after transplanting, and soon after transport to the greenhouse. *Paysonia* plants almost certainly exhibit typical Brassicaceae S-type self-incompatibility (Kitashiba and Nasrallah, 2014), but, because they are predominantly outcrossing, seeds from the same plant can segregate for different S-alleles. We used this property to experimentally cross individuals grown from seed from the same mother plant, and sampled those in which seeds developed normally. This allowed us to reduce heterozygosity between parents, except for the incompatibility alleles. As expected, several crosses between different plants from the same mother consistently failed to produce seed, leading us to assume that these shared the same S-locus. Crosses between plants where seeds developed normally were made reciprocally on a weekly schedule, and unambiguously labeled with jewelers tags to record the male parent and the date of pollination.

After eight weeks of weekly pollinations, we harvested siliques that represented seed development that was between one and eight weeks after pollination. All samples were stored at -80°C until further use.

### 2.2 Developmental staging and morphological analysis

Samples of seeds from each stage were manually isolated from siliques with tweezers on dry ice, and photographed using a S80 APO stereo dissecting microscope with MC120 HD camera (Leica). Up to 10 siliques, with up to 12 seeds per silique, were examined at each stage. Mean seed area, seed length, and seed width were calculated from measurements made in ImageJ (Schneider et al., 2012). Frozen seeds from 14, 21, 28, and 42 days after pollination (DAP) were selected for transcriptome analysis. We chose this sampling regime based on comparison with the developmental timeline presented by Reed et al. (1997), where HFA content started accumulating by 14 DAP and reached a maximum at approximately 35 DAP.

### 2.3 Fatty acid methyl esters (FAMEs) analysis

Approximately 10 mg of each seed sample was mixed with 300 μL of extraction solvent, which consisted of 212 μL of methanol, 60 μL of toluene, 6 μL 2% sulfuric acid, 8μL 0.2% butylated hydroxytoluene, and 15 μL 10 nmol 17:0 TAG as internal standard. The mixtures were homogenized using the bead mill homogenizer at room temperature for 10 minutes at 20 Hz with one 5 mm zirconia bead per sample, and then were incubated for 2 hours at 80° C with 750 RPM in a thermal mixer. After cooling down to room temperature, 300 μL 0.9% NaCl and 300 μL hexane were added to each sample, then vortexed for 20 seconds followed by centrifuging for 3 minutes at 3,000 g, and the resulting supernatants were used for FAMEs analysis.

The quantitative FAMEs analysis was conducted on an Agilent 7890A GC, equipped with a 5975c MS detector with inlet settings: 280 °C, 12.9 psi, total flow 25.2 mL/min, and split ratio 20:1; column settings: SLB-IL111 fused silica capillary column, 1.2 mL/min flowrate, 12.9 psi, constant flow; oven settings: temperature 100 °C hold for 1 min, ramp to 260 °C at 4 °C/min hold for 1 min, ramp to 100 °C at 100 °C/min hold for 4 min, total run time 47.6 min; MS settings: transfer line 260 °C, MS source 230 °C, MS quad 150 °C. Data was processed using Agilent MassHunter Qualitative Analysis software.

An authentic standard of lesquerolic acid methyl ester was purchased from Santa Cruz Biotechnology, while methyl ricinoleate, methyl palmitic acid, methyl heptanoic acid, methyl stearic acid, methyl oleate, methyl linoleate, methyl linolenate, methyl cis-11-eicosenoate, and methyl cis,cis-11,14-eicosenoate were purchased from Sigma Aldrich.

### 2.3 RNA extraction, library preparation and RNA-Seq

RNA extraction of three biological replicates of each stage was performed using the MO BIO PowerPlant RNA isolation kit (Qiagen), according to the manufacturer’s protocol. Each replicate used between 30 and 50 seeds from multiple siliques, with the number depending on the size of the seed at each stage. The quality of extracted RNA was assessed with a 2100 Bioanalyzer (Agilent). The libraries were constructed at the Genomics Core Facility at West Virginia University using the KAPA mRNA Stranded Kit (KAPA Biosystems), and then 150-bp paired-end sequenced at the BioHPC Lab at Cornell University using an Illumina NextSeq 500 platform. All the sequencing files were generated with the Illumina pipeline software v1.8.

### 2.4 *de novo* transcriptome assembly, annotation and differential expression analysis

The raw reads were trimmed and quality-filtered using Trimmomatic v0.36 (Bolger et al., 2014), and then the ribosomal RNA was removed by SortMeRNA v2.1 (Kopylova et al., 2012). The cleaned reads were pooled for *de novo* assembly using the Trinity suite v2.6.6 (Grabherr et al., 2011;Haas et al., 2013) with the default parameters. The results were processed by CD-HIT-EST in order to remove redundant contigs (Li and Godzik, 2006), and the longest contigs were extracted. Gene annotations were predicted with Transdecoder v5.2.0 (http://transdecoder.github.io). The assembled transcripts were annotated with BlastX by searching against the Viridiplantae subset of NCBInr, UniProt and TAIR (The Arabidopsis Information Resource), with an E-value cut-off of 10e-6. The Trinotate v3.1.1 (http://trinotate.github.io) was used to annotate the assembly with Pfam, KEGG SignalP, TmHMM and Gene Ontology.

Transcript abundances were quantified by RSEM (Li and Dewey, 2011), using a built-in script in Trinity, and differentially expressed genes were assessed using edgeR (Robinson et al., 2009), with FDR < 0.05, fold change ≥ 1.5. Genes with an FPKM < 10 were excluded from further analysis. Self-organizing map clustering was performed using MEV v4.8.1 <u>(h</u>t<u>tp://mev.tm4.org/)</u> with the bubble neighborhood method, and radius set to 0.8. GO functional enrichment was performed using topGO package in R, with Fisher’s exact test, and the negative log10 transformed *p*-values were visualized using heatmaps.

### 2.5 Gene cloning and quantitative RT-PCR

cDNA was synthesized with the High Capacity cDNA Reverse Transcription Kit (Applied Biosystems). Primers of *PaFAH12, PaFAD2* and *PaFAD3* and *PaKCS18* were designed based on the contigs recovered from the Trinity assembly and published *Physaria* full-length coding sequences. Sequences were amplified with Q5 or Phusion high-fidelity DNA polymerase (New England Biolabs) from cDNA. The PCR products with expected size were subcloned into pENTR/D-TOPO Vector (Invitrogen), and Sanger-sequenced. For quantitative RT-PCR, standard 2-step amplification reactions were performed using PowerUp SYBR Green Master Mix (Applied Biosystems) with 5 ng cDNA according to the product manual, on a LightCycler 480 System (Roche). Relative transcript abundance was calculated by the ΔCT method using the *Paysonia* 18S rRNA gene as a housekeeping gene.

### 2.6 Phylogenetic analyses

To investigate the evolution of the hydroxylase enzymes in *P. fendleri, P. lindheimeri*, and *Paysonia auriculata*, a phylogeny of *FAD2* and hydroxylase genes from selected Brassicaceae species together with several outgroups was assembled and used to test for evidence of selection in the evolution of the hydroxylases. CDS sequences were downloaded from Genbank, aligned using MUSCLE (Edgar, 2004). Alignments were analyzed in PhyML v3.0 (Guindon et al., 2010), using the General Time Reversible model, with estimated gamma, nucleotide frequencies, and invariant sites, and four gamma rate categories. Topologies were optimized using subtree pruning and rejoining. Bootstrap values were calculated based on 1000 bootstrapped data sets. Alignment and tree files were imported into PAML v4.0 (Yang, 2007), and tested for positive selection on the branch leading to the clade of the hydroxylase sequences. Three separate models were fitted, a branch, a site, and a branch-site model. Amino acid sequences of known enzymes were aligned using MAFFT (Katoh and Standley, 2013), and phylogenetic trees were visualized by iTOL (itol.embl.de).

## 3 Results

### 3.1 Morphological changes during seed development

Seeds from multiple siliques were collected and measured each week for up to 56 DAP. The height and width of each seed was measured using Image J, and seed area was calculated as area within the seed outline (Supplementary Figure 1). Seeds of *Paysonia* go through several morphological changes during development (Figure 1): at 7 DAP the seeds are a pale green color, at 14 DAP the seeds are a translucent dark green color, at 21 DAP the seeds have turned an opaque yellow-green, and after 28 DAP the seeds turn a dark-reddish brown. The seeds reach their maximum size at 21 DAP (Supplementary Figure 1), and then become smaller as they lose water at maturity. The seed number of individual siliques at each stage varied widely, with older siliques having fewer seeds (Supplementary Figure 2), suggestive of seed abortion.

**Figure 1.**
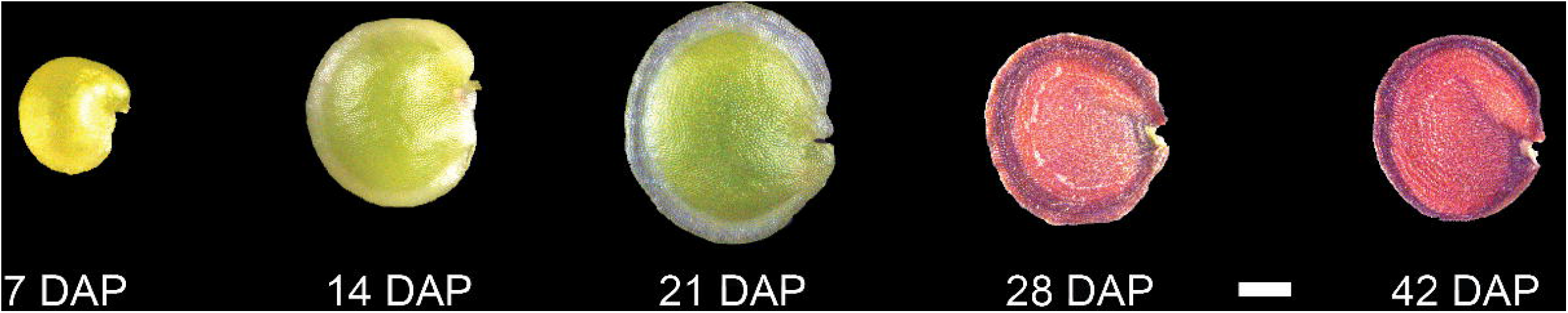
Developmental timeline of *P. auriculata* seed development. Representative seeds from 14, 21, 28, and 42 DAP (days after pollination). Scale bar = 1 mm.

### 3.2 Changes of FAMEs profile

The total lipid content was calculated from the sum of individual FAMEs (Figure 2A). At 14 DAP the relative mean lipid content in the seeds was 2.2%. At 21 DAP, mean lipid content had increased to 4.7%, and then dramatically increased to 20.2% at 28 DAP, and showed little change at 42 DAP with 20.6% mean lipid content.

**Figure 2.**
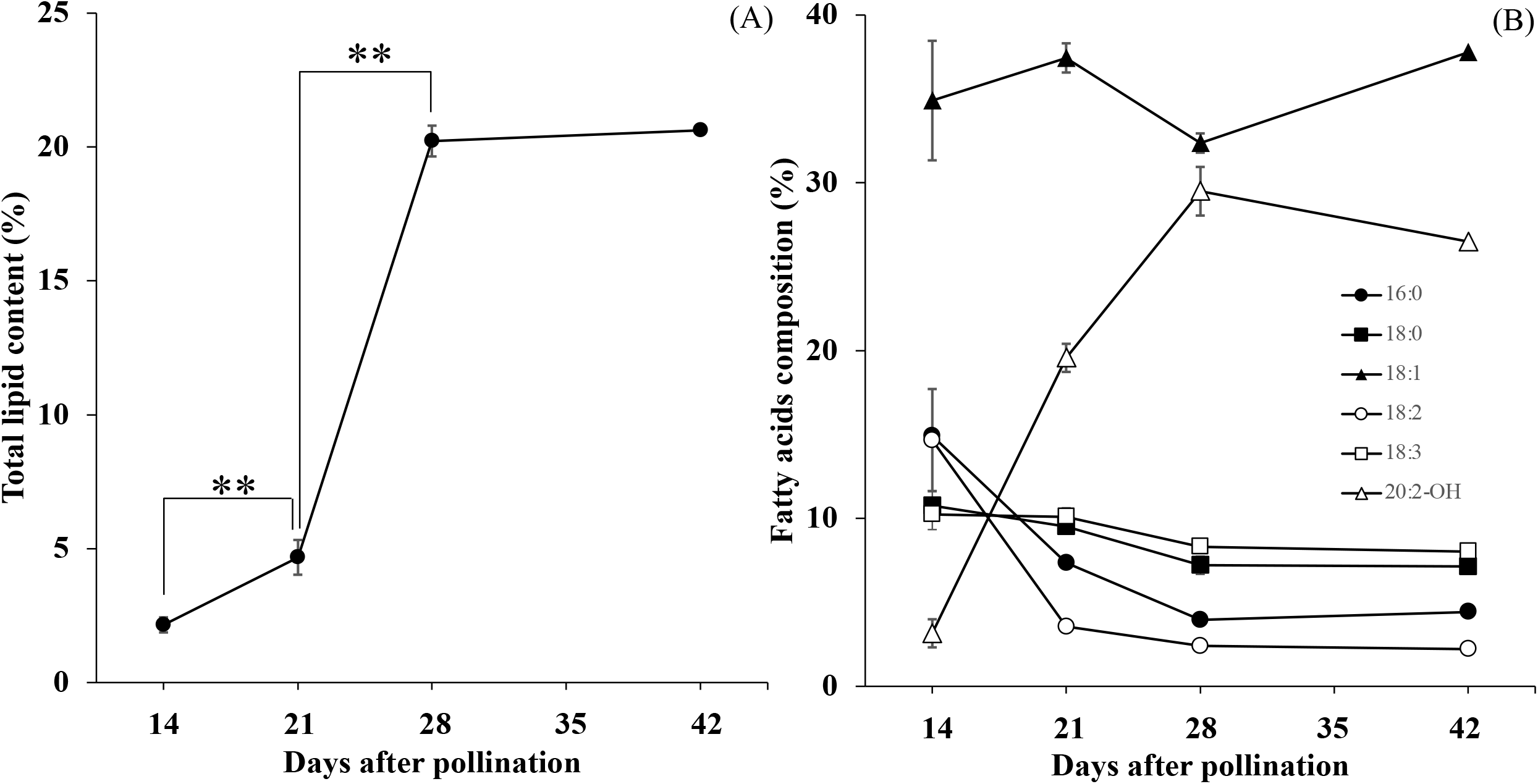
Accumulation of total lipids (A) and six major FA composition (B) during seed development. Each data point represents the mean ± SD of triplicate samples. The double asterisks indicated Significant changes that analyzed by t-test (p < 0.01).

We also measured the relative amounts of each of the components of the total lipids found. We detected 7 FAs and 4 HFAs, and their changes in relative abundance during *Paysonia* seed development is summarized in Table 1. The changes of six major fatty acids (percentage composition ≥ 10% in any stage of seed development) are shown in Figure 2B. At 14 DAP, *Paysonia* seeds had relatively high levels of saturated FAs, with levels of 14.9% for 16:0 and 10.8% for 18:0. Unsaturated FAs had similar levels except for 18:1, which was at almost 35%. The HFAs were at low levels, with 2.7% for 18:1-OH, 3.8% for 20:1-OH, and 3.2% for 20:2-OH. At 21 DAP, there was an overall decrease in saturated and unsaturated FAs, except for 18:1 OH, which climbed slightly to 37.4%. The HFA contents were also similar to 14 DAP, except for a sharp increase in 20:2-OH from 3.2% to 19.6%. At 28 DAP saturated and unsaturated FAs maintained their levels, but there was a slight rise in 20:1-OH to 6.7 % and a larger rise in 20:2-OH to 29.5%. At 42 DAP there was relatively little change in percentages from those seen at 28 DAP.

**Table 1.** The fatty acid composition of total lipid in developing seeds 14, 21, 28, and 42 days after pollination (DAP). Each FA fraction was measured as the percentage of total FA. Each value is presented as the mean ± SD of triplicate samples. 16:0 is palmitic acid; 16:1 is palmitoleic acid; 18:0 is stearic acid; 18:1 is oleic acid; 18:2 is linoleic acid; 18:3 is linolenic acid; 20:1 is eicosenoic acid; 18:1-OH is ricinoleic acid; 18:2-OH is densipolic acid; 20:1-OH is lesquerolic acid; 20:2-OH is auricolic acid.

| Fatty Acid<br>(Mol %) | 14 DAP | 21 DAP | 28 DAP | 42 DAP |
| --- | --- | --- | --- | --- |
| 16:0 | 14.9 ± 0.8 | 7.4 ± 0.3 | 4 ± 0.1 | 4.4 ± 0.2 |
| 16:1 | 3.4 ± 0.3 | 2.6 ± 0.3 | 1.4 ± 0.1 | 1.5 ± 0.1 |
| 18:0 | 10.8 ± 0.9 | 9.5 ± 0.4 | 7.2 ± 0.5 | 7.1 ± 0.2 |
| 18:1 | 34.9 ± 3.6 | 37.4 ± 0.9 | 32.4 ± 0.6 | 37.8 ± 1.8 |
| 18:2 | 14.7 ± 3.0 | 3.6 ± 0.1 | 2.4 ± 0.2 | 2.2 ± 0.1 |
| 18:3 | 10.2 ± 0.9 | 10.1 ± 0.5 | 8.3 ± 0.1 | 8 ± 0.3 |
| 20:1 | 1.5 ± 0.5 | 2.8 ± 0.2 | 3.1 ± 0.4 | 2.8 ± 0.3 |
| 18:1-OH | 2.7 ± 0.7 | 2.7 ± 0.3 | 3.9 ± 0.1 | 3.7 ± 0.2 |
| 18:2-OH | 0.0 ± 0.0 | 0.7 ± 0.1 | 1.2 ± 0.1 | 1.5 ± 0.2 |
| 20:1-OH | 3.8 ± 1.1 | 3.6 ± 0.5 | 6.7 ± 0.7 | 4.4 ± 0.7 |
| 20:2-OH | 3.2 ± 0.8 | 19.6 ± 0.8 | 29.5 ± 1.4 | 26.5 ± 1.3 |

### 3.2 RNA-Seq and *de novo* transcriptome assembly

After filtering low quality reads and rRNA sequences, 31,471,350 to 41,054,622 Ilumina reads were retained in each library (Supplementary Table 1). The raw data are deposited in the Short Read Archive (SRA) database of NCBI under accession number PRJNA630887. The Trinity suite produced a *de novo* transcriptome assembly with 226,807,152 bases, which represents 270,114 contigs with average length of 840 bp and N50 length of 1,255 bp. These contigs represent the putative transcripts that were independently assembled in our 12 sequencing libraries. To reduce complexity of the assembled sequences, 122,730 of the longest contigs were extracted from the assembly. Among them, more than 55% (677,110) were within 200-400 bp, ∼35% (43,295) were within 400-1,500 bp, and ∼9.5% (11,725) were longer than 1,500 bp. The detailed assembly statistics are in Supplementary Table 2. Although the De Bruijn Graphs-based Trinity has been proven to be a powerful approach for assembling *de novo* transcriptome in the absence of a reference genome, it still has some limitations, and may yield more redundant contigs than the actual number of genes expressed (Haas et al., 2013;Sánchez-Sevilla et al., 2017). These contigs may originate from different alternative splicing variants or paralogous transcripts, and cannot be further validated in the absence of a complete genomic sequence.

The quality of the *de novo* assembly was further evaluated by realignment of the cleaned reads to the 122,730 longest contigs, by which 55.98-63.92% reads were uniquely aligned to the assembly, 4.19-7.36% reads were aligned to more than one location on assembly, and the overall alignment rate was 77.88-83.25% (Supplementary Table 1).

To annotate the transcriptome assembly, we performed BlastX against several annotation databases with an e-value cutoff of 1e-5. The annotation ratios with at least one hit were 48% (58,328), 32% (39,597) and 42% (51,607) in NCBInr, UniProt and TAIR, respectively. Additionally, 20% (24,858) were identified in Pfam, and 28% (34,515) were annotated in the KEGG database (Supplementary File 1).

Gene ontology (GO) analysis was used to functionally categorize *Paysonia* contigs. A total of 32% (39,723) contigs were assigned with at least one GO term. Of these, 84% (33,519) of the annotated contigs were assigned to Biological Process, 84% (33,525) to Cellular Component, and 86% (34,207) to Molecular Function. The terms “cellular process” (27,501), “metabolic process” (24,528) and “response to stimulus” (12,375) were prominently represented in the Biological Process category. The terms “cell” (31,329), “cell part” (31,199) and “organelle” (25,037) were the most highly represented groups in the Cellular Component category. “Binding” (25,941), “catalytic activity” (21,522) and “transcription regulator activity” (2,895) were the top three terms in the Molecular Function category.

The Arabidopsis Acyl-Lipid Metabolism Database (AraLip, http://aralip.plantbiology.msu.edu/) (Li-Beisson et al., 2013), which represents a curated collection of genome-wide lipid genes in Arabidopsis, was used to mine fatty acid metabolic genes from our reference assembly. A total of 766 non-redundant Arabidopsis lipid-related genes from 16 sub acyl-lipid functional groups were used to query the *Paysonia* reference assembly with BlastN. We found homologs of approximately 83% (643 out of 766) of Arabidopsis lipid genes, with sequence identities of 74-97% and a maximal e-value of 2.8e-08, including 93% of FA and 81% of TAG synthesis genes. This suggests that our transcriptome is highly representative of fatty acid and TAG biosynthesis genes in *P. auriculata*.

### 3.3 Differentially expressed genes

Differentially expressed genes (DEGs) were assessed by pairwise comparisons of the expression levels of the longest contigs (here referred to as gene expression levels) between developmental stages. In a comparison between 21 DAP and 14 DAP, 5,154 DEGs were detected, with 2,036 being up-regulated and 3,118 down-regulated. In a comparison between 28 DAP and 21 DAP, 3,893 DEGs were identified, with 1,296 being up-regulated and 2,597 down-regulated. In a comparison between 42 DAP and 28 DAP, only 5 DEGs were found, and they all were down-regulated. Overall, a total of 7,869 unique DEGs were identified from all three pairwise comparisons (Supplementary File 1).

Self-organizing map (SOM) clustering was used to assess global transcriptome dynamics and patterns during seed development (Figure 3). All 7,869 DEGs could be assigned into four distinct clusters (Supplementary File 1). Cluster 1 contained the most genes (4,396), and had peak expression at 14 DAP. Cluster 2 was upregulated at 28 and 42 DAP, cluster 3 was upregulated at 21 DAP and then reduced at 28 and 42 DAP, and cluster 4 was upregulated at 21 and 28 DAP and down-regulated at 42 DAP. A GO enrichment analysis was performed to evaluate the functions of genes in each cluster (Supplementary Table 3). Genes related to photosynthesis, and carbohydrate and fatty acid biosynthesis were especially well-represented in cluster 1. Genes related to lipid catabolic process, lipid modification, and oxidative stress were enriched in cluster 2. The genes in cluster 3 were mostly annotated as phenylpropanoid metabolism and suberin biosynthesis genes. In cluster 4, genes were mostly related to sucrose metabolism, response to water deprivation, aging and lipid localization.

**Figure 3.**
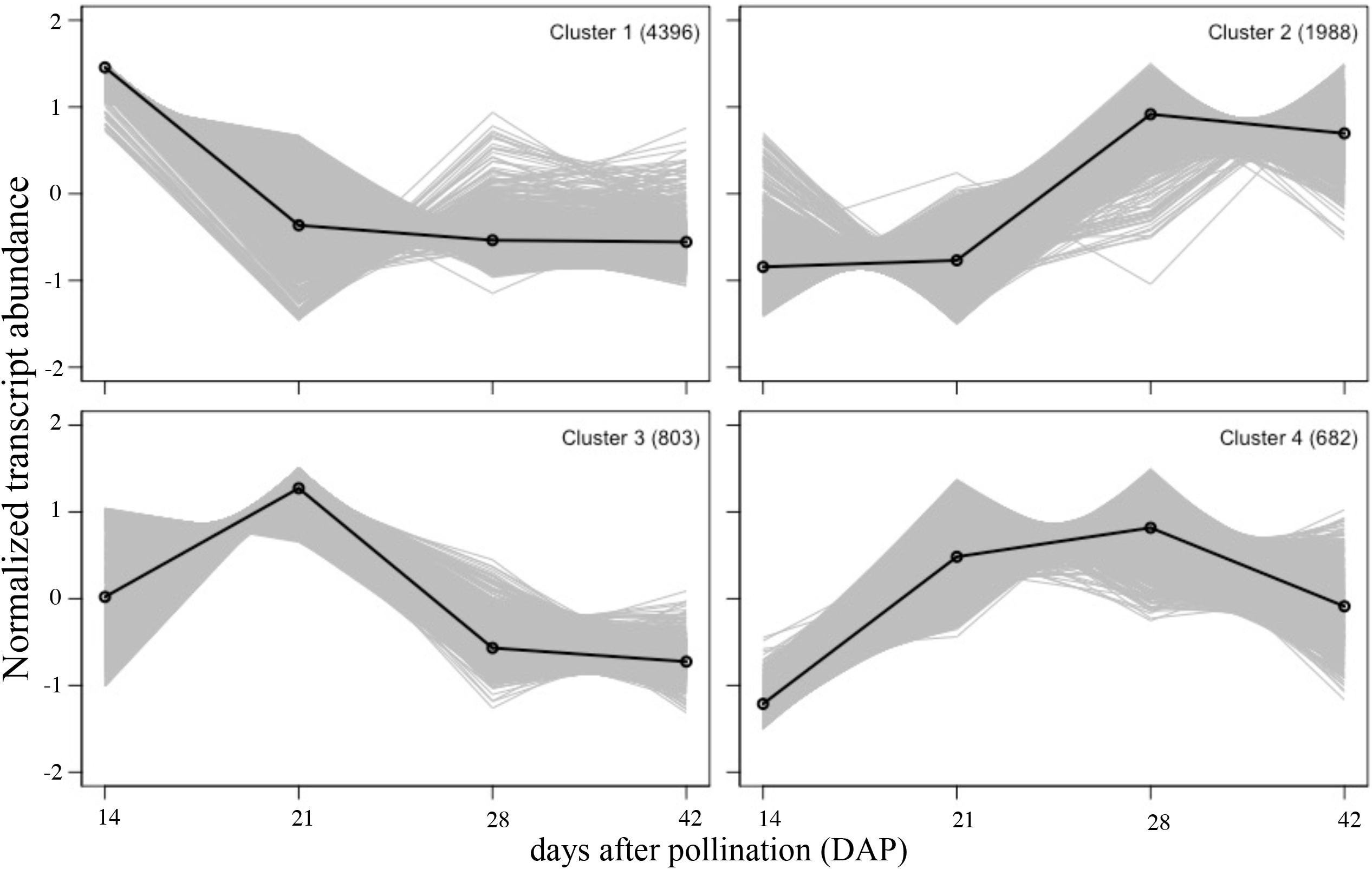
Self-organizing map (SOM) clustering. SOM clusters of DE genes showing their expression patterns during four stages of seed development. The number of DE genes in each cluster is indicated in top right. Solid lines represent the group mean of gene expressions in cluster. Each stage is represented by three biological replicates.

### 3.4 Differential expression of transcription factors regulating seed development and FA biosynthesis

In Arabidopsis, *LEAFY COTYLEDON 1* (*LEC1*), along with *ABSCISIC ACID INSENSITIVE3* (*ABI3*), *FUSCA3* (*FUS3*) and *LEAFY COTYLEDON 2* (*LEC2*), are known to be the master regulators of seed development and seed maturation, such as storage protein/FA/TAG synthesis, and desiccation (Sreenivasulu and Wobus, 2013). *WRINKLED1* (*WRI1*) is hypothesized to be the direct regulator of FA biosynthesis (Focks and Benning, 1998;Cernac and Benning, 2004;Baud et al., 2009) and glycolysis, under the control of *LEC1* (Sreenivasulu and Wobus, 2013). In our study, only one *ABI3* contig was found (Figure 4 and Supplementary File 1), peaking at 21 DAP (in cluster 3 of SOM), while the contigs of *LEC1, LEC2* and *FUS3* were weakly expressed (average FPKM<10) and did not show differential expression. Four contigs of *WRI1* exhibited high transcript abundance at 14 DAP (Figure 4 and Supplementary File 1), then gradually decreased (in cluster 1 of SOM) (Figure 3). The expression patterns of *ABI3* and *WRI1* were similar to those found in the expression analyses in *Physaria* (Horn et al., 2016).

**Figure 4.**
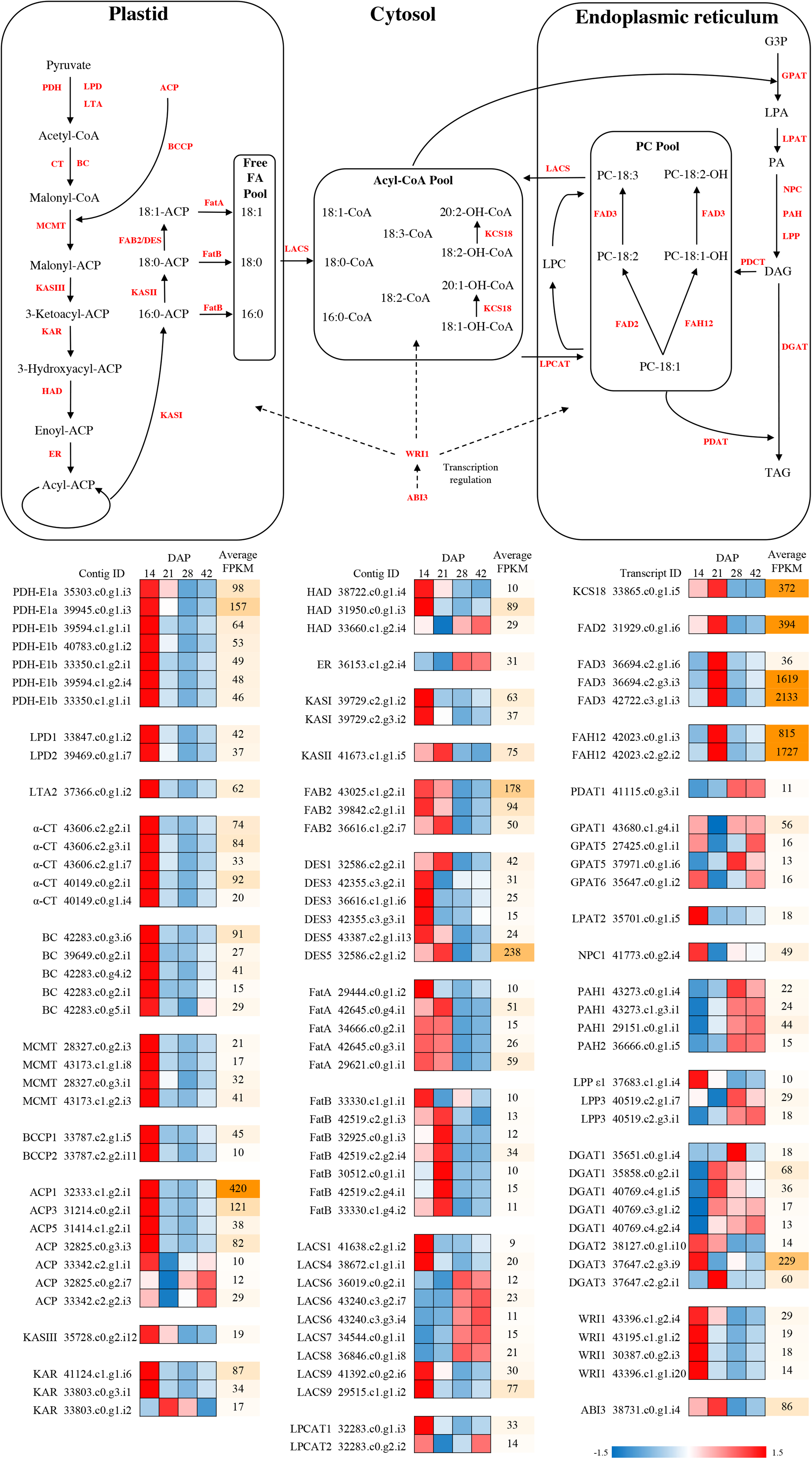
Fatty acid synthesis pathways with differential expression during seed development in *P. auriculata*. The differential expressed genes are presented with mini heatmaps that exhibit their normalized transcript abundance at four stages (from left to right: 14, 21, 28, and 42 DAP), the genes without differential expression are indicated as gray fonts. The dashed lines indicated the transcription regulation in FA/HFA biosynthesis. Abbreviations: ACP, acyl carrier protein; BC, biotin carboxylase subunit of acetyl-CoA carboxylase; BCCP, biotin carboxyl carrier protein subunit of acetyl-CoA carboxylase; CT, carboxyltransferase subunit of acetyl-CoA carboxylase; DGAT, diacylglycerol acyltransferase; ER, enoyl-ACP reductase; FAB2/SAD, stearoyl-ACP desaturase; FatA, acyl-ACP thioesterase A; FatB, acyl-ACP thioesterase B; FAD2, Δ12 oleic acid desaturase; FAD3, Δ15 linoleic acid desaturase; FAH12, Δ12 oleic acid hydroxylase; GPAT, glycerol-3-phosphate acyltransferase; HAD, hydroxyacyl-ACP dehydrase; KAR, ketoacyl-ACP reductase; KASI, ketoacyl-ACP synthase I; KASII, ketoacyl-ACP synthase II; KASIII, ketoacyl-ACP synthase III; KCS18, 3-ketoacyl-CoA synthase 18; LACS, long-chain acyl-CoA synthetase; LPD, dihydrolipoamide dehydrogenase; LPAT, lysophosphatidic acid acyltransferase; LTA, dihydrolipoamide acetyltransferase; MCMT, malonly-CoA ACP malonyltransferase; PDAT, phospholipid diacylglycerol acyltransferase; PDCT, phosphatidylcholine diacylglycerol cholinephosphotransferase; PDH, pyruvate dehydrogenase; PPase, phosphatidate phosphatases; ABI3, ABSCISIC ACID INSENSITIVE 3; LEC1, LEAFY COTYLEDON 1; WRI1, WRINKLED 1; G3P, glycerol-3-phosphate; LPA, lysophosphatidic acid; PA, phosphatidic acid; DAG, diacylglycerol; TAG, triacylglycerol; LPC, lysophosphatidylcholine; PC, phosphatidylcholine.

### 3.5 Differential expression of FA-related genes

Plastidial *de novo* FA synthesis is the initial step in synthesizing precursors for both common FAs and HFAs, and is regulated by approximately 29 genes (Li-Beisson et al., 2013;Horn et al., 2016). This stage can be divided into two major steps: 1) FA synthesis from pyruvate, to produce common C16:0 FAs, and 2) elongation and desaturation of C16:0 FAs to produce C18:0 and C18:1 FAs, which are finally exported as CoA from the plastid. We detected 20 DEGs in the FA synthesis step (Figure 4 and Supplementary File 1), all of which grouped into Cluster 1 in the SOM analysis, and all of these genes showed highly upregulated expression at 14 DAP followed by a decrease at 21 and 28 DAP. Additionally, 14 DEGs were found in the FA elongation, desaturation and export step (Figure 4 and Supplementary File 1), and their overall patterns were similar to the FA synthesis genes, and were thus also detected in Cluster 1. However, even though some desaturases (FatA/Bs) and thioesterases (FAB2s) were grouped into cluster 1, they showed a somewhat different expression pattern, being upregulated at 14 DAP and remaining upregulated at 21 DAP, before being reduced at 28 and 42 DAP.

The C16/18-CoA produced from FA synthesis is imported into the ER-associated Kennedy pathway, and then acylated twice with *GPAT* and *LPAT*, to yield phosphatidic acid (PA) (Haslam et al., 2016). We detected all nine contigs of *GPAT*, but only *GPAT4, GPAT5*, and *GPAT6* showed elevated expression at 21 DAP (in cluster 3 of SOM) (Figure 4 and Supplementary File 1). Four of five contigs of *LPAT* (*LPAT2, LPAT3, LPAT4* and *LPAT5*) were identified (Figure 4 and Supplementary File 1), all of which showed a trend of downregulation as the seeds matured, even though the variation between replicates rendered the trend non-significant. The PA is hydrolyzed by phosphatidate phosphatases (PPases) to form DAG, which is then used as a substrate for producing TAG by *DGAT* (Li-Beisson et al., 2013). Three PPases, including lipid phosphate phosphatases (*LPPs*), phosphatidic acid phosphohydrolase (*PAHs*) and non-specific phospholipase C (*NPCs*), were found (Figure 4 and Supplementary File 1). Both *PAH1* and *LPP3* were significantly upregulated at 28 and 42 DAP (in cluster 2 of SOM), while *NPC2* and *NPC6* had peak expression at 14 DAP (in cluster 1 of SOM), and *NPC4* was exclusively upregulated at 21 DAP (in cluster 3 of SOM). All three contigs of *DGAT* were detected (Figure 4 and Supplementary File 1), with *DGAT1* being highly induced at both 21 and 28 DAP (in cluster 4 of SOM), *DGAT2* was not differentially expressed, and *DGAT3* was upregulated only at 21 DAP (in cluster 3 of SOM).

The PC pool is the major substrate for FA modifications, including hydroxylation and desaturation, and is sustained by two known routes: 1) acyl-editing by *LPCAT*, and 2) symmetrical head-groups exchange of DAG and PC by *PDCT*. We detected all five PC-channeling enzymes (Figure 4 and Supplementary File 1), *LPCAT1* and *LPCAT2, PDAT1, PDAT2* and *PDCT*, but only *LPCAT1* was significantly upregulated at 14 DAP (in cluster 1 of SOM) (*PDAT1* has a similar pattern, but did not pass the DE significance cutoff). The resulting PC-18:1 can be either hydroxylated to PC-18:1-OH by *FAH12* followed by desaturation by *FAD2*, or directly desaturated to PC-18:2 by *FAD3*. All four HFA modification-related genes were identified (Figure 4 and Supplementary File 1), with three of them, *FAH12, FAD2*, and *FAD3* being highly upregulated at 21 DAP (in cluster 3 of SOM). Both *FAH12* and *FAD3* were much more highly expressed than *FAD2. KCS18* was only induced, at relatively low levels, at 14 DAP (in cluster 1 of SOM) (Figure 4 and Supplementary File 1).

### 3.6 Characterization of FAH12, FAD2, FAD3 and KCS18 in *Paysonia*

In order to obtain further insight into the function of four key HFA genes, *FAH12, FAD2, FAD3* and *KCS18*, we amplified and cloned full-length coding sequences, using primers based on the contigs recovered from our *Paysonia de novo* assembly and published *Physaria* sequences (primers used are listed in Supplementary Table 4). The full-length coding sequences for each gene (GenBank ID: MT496774-MT496777) were successfully verified by Sanger sequencing, and the translated protein sequence alignment showed that *P. auriculata* PaFAH12 had exactly the same AA length (384 AA) as *P. fendleri* PfFAH12. PaFAH12 had high sequence similarity to PfFAH12 (95.3%) and PlFAH12 (95.1%). Emphasizing the close relationship between desaturases and hydroxylases, PaFAH12 also had comparable sequence similarity to PaFAD2 (94.3%) and PfFAD2 (93.5%). The sequence similarity of desaturases (FAD2s and FAD3s), and 3-ketoacyl-CoA synthases (KCSs)were also high (90.1-99.5%) (see Supplementary Figures 3-8 for phylogenetic trees and amino acid alignments, and Supplementary File 2 for percent of sequence similarities of FAH12, FAD2, FAD3 and KCS18 in various species).

We further employed real-time qPCR (RT-qPCR) to assess the accuracy of the transcriptome quantification for the four FA modification genes. Overall, the RT-qPCR expression patterns of *FAH12, FAD2, FAD3* and *KCS18* were consistent with the RNA-Seq results (Figure 5), although some levels at some stages differed (e.g. the wide, though non-significant, difference between the qPCR and RNAseq results at 21 DAP for *FAD2*). The addition of an additional earlier stage (7 DAP) in the qPCR analyses allowed us to confirm that high expression peaks for *FAD2* and *KCS18* do in fact represent the time of maximum upregulation. Similarly, the peak in expression at 21 DAP for *FAH12* and *FAD3* indicate that these do appear to be upregulated later in development than *FAD2* and *KCS18*.

**Figure 5.**
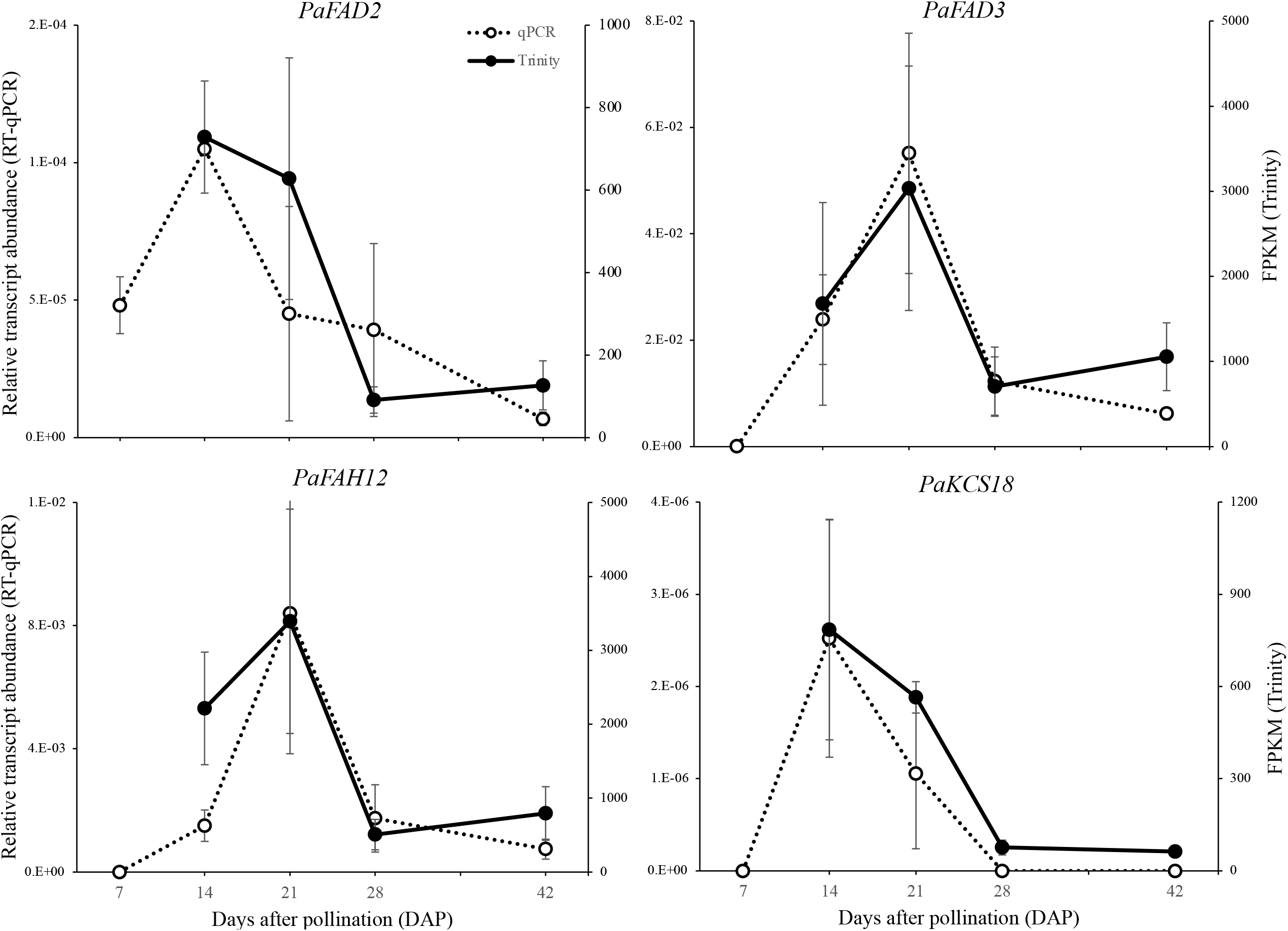
Comparison of the expression of HFA modification-related genes between the transcriptomic and quantitative PCR methodologies. The left and right axis represent the expression level quantified with qRT-PCR (relative transcript abundance) and Trinity (FPKM), respectively. Data are meansL±LSD of nL=L3 biological replicates.

### 3.7 Phylogenetic analysis of *PaFAD2* and *PaFAH12*

A phylogeny of *FAD2* and *FAH12* from selected Brassicaceae species and outgroups was used to test for evidence of selection in the branch leading to the *Paysonia* and *Physaria* hydroxylase sequences (Supplementary Figure 9). Tests for variation in ω (dN/dS) between the branch leading to the *Paysonia* and *Physaria* hydroxylase sequences and the rest of the tree showed a significant difference in ω (*p* < 0.05). A branch-sites model was tested to analyze whether positive selection could be identified for particular amino acids, as a mean difference in ω could either be the product of positive selection or of relaxed purifying selection. The branch-sites model was non-significant, implying that the changed ω on the branch leading to the hydroxylase sequences was due to a relaxation of the strong purifying selection on the branches in the rest of the tree (ω = 0.09 for background branches, ω = 0.18 for hydroxylase branch).

The close relationship between desaturases and hydroxylases has been noted previously (Broun et al., 1998a;Shanklin and Cahoon, 1998), and we decided to combine the data derived from this study with previous published sequences to explore potential conserved hydroxylase-specific residues. We retrieved all published FAH12 sequences from GenBank, including five from plants (*P. lindheimeri, P. fendleri, Orychophragmus violaceus, R. communis* and *Hiptage benghalensis*) and one from fungus *Claviceps purpurea*, and aligned them with the *PaFAH12* sequence and with sequences of *FAD2* from these and other species (Figure 6). We found two residues that were strictly conserved in each type of enzyme, G105 and F218 in all *FAH12s*, and A105 and Y218 in all *FAD2s*, except in fungal *CpFAD2*. The residue G105A right before the first histidine box (HECGHx) was previously suggested as a critical residue to identify hydroxylases from desaturases (Cahoon and Kinney, 2005). T148 was found to be conserved in all *FAD2s*, but divergent in *FAH12s*. In addition, the motif right after the third histidine box (HVAHH) is highly conserved (LFSTMP) in all plant *FAD2s*, but were divergent in *FAH12s*. Interestingly, *PaFAH12* only has one residue that was different from the LFSTMP motif (S322A), whereas other *FAH12s* had two or more differences. Otherwise, all plant *FAH12* and *FAD2*, including *PaFAH12* and *PaFAD2*, possessed an ER localization signal motif (Φ-X-X-K/R/D/E-Φ) (McCartney et al., 2004).

**Figure 6.**
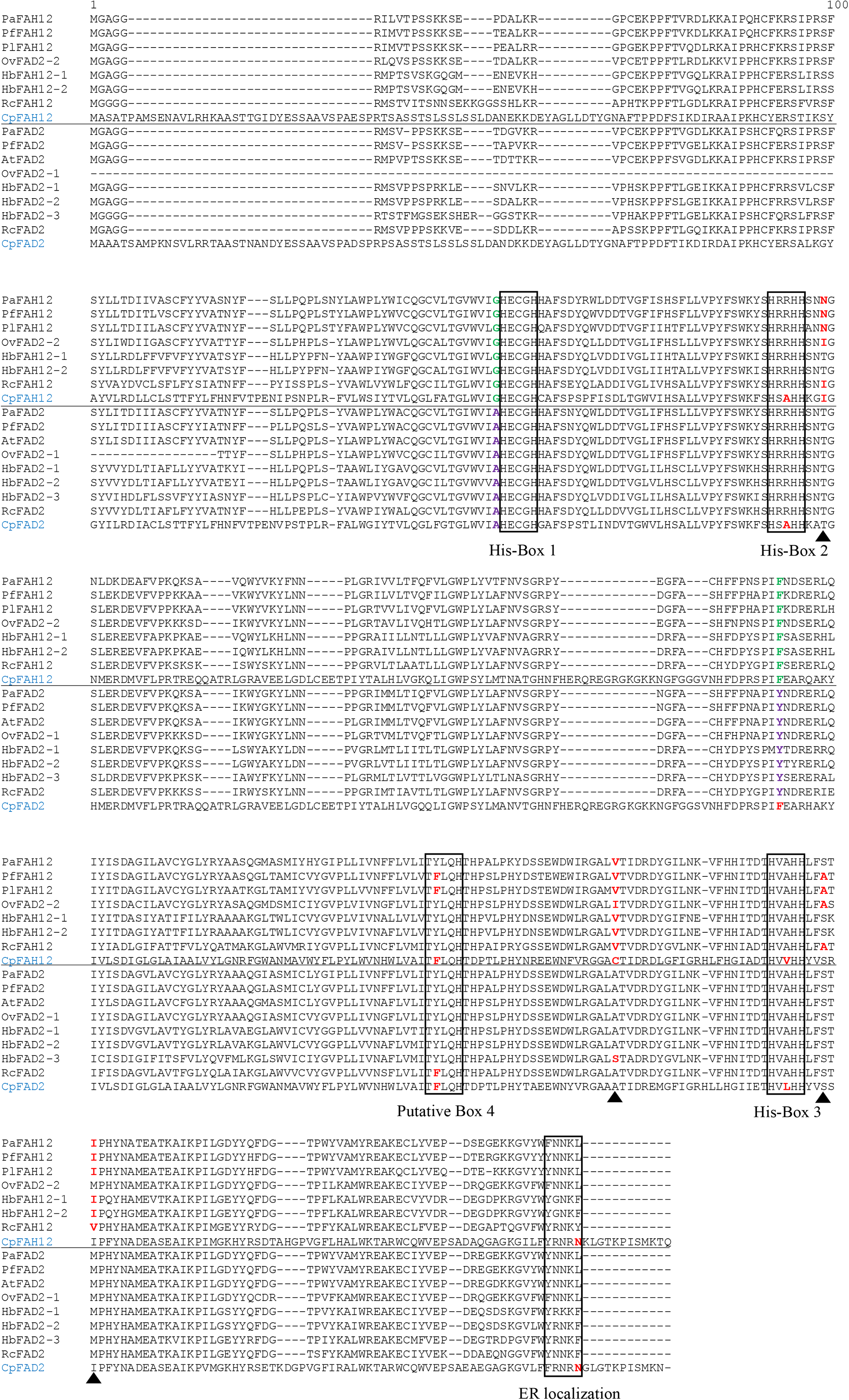
Alignment of PaFAH12 and PaFAD2 with other published FAH12s and FAD2s. Species included *P. fendleri, P. lindheimeri, A. thaliana, O. violaceus, H. benghalensis, R. communis* and fungus *C. purpurea*. Please note, OvFAD2-2 is characterized as a hydroxylase (Li et al., 2018). The putative box 4 is hypothesized by Robin et al. (Robin et al., 2019). The ER localization signal motif (ΦLXLXLK/R/D/ELΦ) is proposed by McCartney et al. (McCartney et al., 2004), with the exceptions of fungus CpFAH12 and CpFAD2, where the CLterminal Φ is a hydrophilic amino acid (Asn) but not a hydrophobic amino acid. The unconsensus residues within sequence of interest were red-bolded. The residues that were strictly conserved in each type of enzyme (A105 and Y218 in all FAD2s, except in fungus CpFAD2) were green and purple-bolded. The four essential residues (T148N, A296V, S322A, M324I) that converted AtFAD2 into a hydroxylase (Broun et al., 1998b) were labeled with black triangles. Sequences used in this alignment: PfFAH12 (AAC32755), PlFAH12 (ABQ01458), OvFAD2-2 (APQ40636), HbFAH12-1 (AGN95845), HbFAH12-2 (AGN95846), RcFAH12 (XP_002528127), CpFAH12 (BAW27658), PfFAD2 (AFP21683), AtFAD2 (NP_001319529), OvFAD2-1 (APQ40635), HbFAD2-1 (AGN95841), HbFAD2-2 (AGN95842), HbFAD2-3 (AGN95843), RcFAD2 (NP_001310648), CpFAD2 (ABS18716).

## 4 Discussion

In this study, we investigated the time-course of seed development in *P. auriculata*, analyzing morphological, fatty acid composition and gene expression changes. We applied RNA-seq and *de novo* transcript assembly to identify orthologs and expression patterns of almost all fatty acid synthesis and modification genes, including the gene encoding the hydroxylase enzyme.

The timing of the various stages of seed development in *P. auriculata* appears to be somewhat faster than in *Physaria*, with the characteristic change of the seed coat color from green to brown as the seed matures happening between 21 to 28 days in *P. auriculata*, as opposed to between 35 and 42 days in *P. fendleri* (Chen et al., 2009), and 42 to 49 days in *P. lindheimeri* (Chen et al., 2017). The approximate timing of maximum seed size in *P. auriculata* is at 21 DAP whereas in *P. fendleri* and *P. lindheimeri* it is at approximately 35 DAP. In addition, the *P. auriculata* seed is approximately 3 times as large as the *Physaria* seed, suggesting that its rate of growth is much faster than seeds of *Physaria*. As with *Physaria*, seeds increase in size up until the start of seed maturation, after which seeds begin to shrink due to water loss. Seed numbers per silique are less than those reported for *Physaria*, with a mean of less than 10 at maturity as opposed to over 16 in *P. fendleri* (Brahim et al., 1998). Therefore, the seeds of *P. auriculata* appear to develop faster and attain a larger size than in *Physaria*, but have fewer seeds per silique at maturity. Often there are aborted seeds along with viable seeds within a pod, suggesting that there may be competition for nutrients in the developing siliques.

*Paysonia auriculata* seeds had only 2.2% total oil at the earliest measurement at 14 DAP, with approximately 25% saturated FAs (16:0 and 18:0), and 65% unsaturated FAs (16:1, 18:1, 18:2, 18:3 and 20:1), similar in amount and proportion to the two *Physaria* species (Chen et al., 2011;Chen et al., 2017). *P*.*auriculata* has its highest oil content at maturity (20.6% at 42 DAP), which is slightly lower than *P. fendleri* (27.6% at 49 DAP) (Chen et al., 2009), but much higher than *P. lindheimeri* (10.9% at 56 DAP) (Chen et al., 2017). However, the HFA content at maturity is lower in *P. auriculata* (35.6%) than either *P. fendleri* (77.7%) (Chen et al., 2011) or *P. lindheimeri* (59.5%) (Chen et al., 2017). One interpretation of these differences is that only one of the three fatty acids in *P. auriculata* TAG is an HFA, whereas on average, two of the three are in the two *Physaria* species. The greatest rate of production of the major HFA in all three species was at approximately similar times: 20:2-OH in *P. auriculata* between 21 and 28 DAP, and 20:1-OH in *Physaria fendleri* and *P. lindheimeri* between 21 and 28 DAP (Chen et al., 2011;Chen et al., 2017). The accumulation of 20:2-OH HFA in *P. auriculata* also coincided with the upregulation of *PaFAH12, PaFAD3* and PaKCS18 expression, as was also reported in *P. fendleri* (Chen et al., 2011;Kim and Chen, 2015;Horn et al., 2016) and *P. lindheimeri* (Chen et al., 2021). Both *P. auriculata* and *P. fendleri* maintain relatively high and constant levels of 18:1 unsaturated FA during seed development (32.4-37.8% and 28.2-37.0% respectively), whereas *P. lindheimeri* is reported to have lower levels that peak when HFA production first accelerates at 19.3% and then fall in subsequent sampling periods to 5.6%. (Chen et al., 2011;Chen et al., 2017).

The RNAseq analyses of seed development over the four sampling stages succeeded in identifying orthologs of almost all of the FA synthesis and modification genes that are described in the Arabidopsis acyl-lipid metabolism database, and that we would expect from the published transcriptomes of *P. fendleri* and *P. lindheimeri*. Therefore, our analyses have likely captured a comprehensive representation of seed development genes in *P. auriculata*. The differentially expressed gene analysis showed that changes from 14 to 28 DAP make up the bulk of the growth phase of the seed, with little change between 28 and 42 DAP. We restricted our analyses to the longest contigs in this study, but recognized that other shorter contigs may represent paralogous copies, alternatively spliced transcripts, or redundant transcripts from different libraries. Distinguishing these will require a future deep sequencing of the whole genome. The GO analyses indicated that the overall stages of embryogenesis, growth and maturation align well with the characterization of the transcriptomes of *P. fendleri* and *P. lindheimeri*.

The genes involved in FA synthesis and elongation were generally found in Cluster 1 and were highly upregulated in early development (14 DAP), and then gradually decreased later in development (28 DAP and later). On the other hand, most of the HFA and Kennedy pathway-related genes displayed a delayed response, being upregulated in the mid stage (21 and/or 28 DAP, Cluster 3), with the exception of *DGAT*, the enzyme catalyzing the final and rate-limiting step of TAG synthesis, which was dramatically upregulated at the late maturation stage (42 DAP, Cluster 2). The order of these patterns of gene upregulation are similar to that previously reported for *Physaria* (Kim and Chen, 2015;Horn et al., 2016), although the highest expression of PaFAH12 we found was at 21 DAP in *Paysonia*, rather than ∼30 DAP in *Physaria* (Horn et al., 2016).

The PC pool as the key substrate for HFA production is hypothesized to have two inputs in the form of *PDCT* and *LPCAT*, and one output connecting DAG and TAG by *PDAT* (Bates and Browse, 2012). In our analyses, *LPCAT* is the only differentially expressed gene among them, peaking at 14 DAP and declining thereafter. *LPCAT* encodes the enzyme that is responsible for importing newly synthesized FAs from the acyl-CoA pool to the PC pool. Our results are at too coarse a scale to track the order of biosynthetic steps with any precision, but may suggest that in *Paysonia LPCAT*-mediated acyl editing is the major driving force to channel common FAs into the PC pool. The hydroxylated PC (18:1/18:2-OH) can be exported by PPases to the cytosolic Acyl-CoA pool, to be further elongated to hydroxylated CoA (20:1/20:2-OH-CoA), or alternatively, the HFA-PC can be directly incorporated into DAG and form HFA-TAG, by *PDAT* (Li-Beisson et al., 2013).

When the main FA synthesis and modification genes are considered, the export of HFAs from the plastid mediated by *LACS9*, the desaturation of 18:0 to 18:1 by *FAD2*, the hydroxylation of 18:1 to 18:1-OH by *FAH12*, and the elongation of 18:1-OH to 20:1-OH by *KSC18* all occur earlier in the development of *P. auriculata* seeds than in *P. fendleri*. The processes mediated by enzymes such as *FAD3*, catalyzing the desaturation of 20:1-OH to 20:2-OH, and *DGAT1* and *DGAT3* that create TAG appear to act at a similar stage in the two species. In our transcriptome data, we found that *FAD3* expression peaked at 14 DAP, and it was the most abundant transcript throughout seed development (average FPKM = 2133 and 1619 of two contigs). This fact is consistent with the role of *FAD3* in desaturation of 18:1-OH and 20:1-OH, and suggests its importance in generating 20:2-OH, as suggested previously (Broun and Somerville, 1997;Reed et al., 1997). Studies in *Physaria* report two transcripts of *FAD3* (*PfFAD3*-1 and *PfFAD3*-*2*), with distinct seed expression patterns and protein sequences (Li-Beisson et al., 2013;Kim and Chen, 2015;Lee et al., 2015). However, in our study, we find only one *PaFAD3* transcript, whose sequence was very similar to PfFAD3-1 (95%), but not to PfFAD3-2 (78%). The peak of *PaFAD3* at 21 DAP agrees with the reported *PfFAD3* expression pattern (Horn et al., 2016).

Of the four HFA-related genes, *PaFAD2* and *PaKCS18* both peaked in expression at 14 DAP while *PaFAD3* and *PaFAH12* peaked at 21 DAP and had relatively low expression at 14 DAP. *FAH12* is the key gene encoding oleate 12-hydroxylase, whose function is to hydroxylate PC-18:1-PC to PC-18:1-OH. Broun et al. (1998b) reported an amino acid substitution assay, which successfully converted Arabidopsis *AtFAD2* into a hydroxylase by replacing four amino acids (T148N, A296V, S322A, M324I) that were adjacent to histidine boxes with *PfFAH12* residues, but they did not find any single amino acid that could determine the hydroxylation activity in an additional reciprocal replacement assay. Interestingly, a recent chimera and mutagenesis analysis of fungus *CpFAH12* (Robin et al., 2019) identified the isoleucine at 198 (I198) as likely the single crucial residue that is responsible for the hydroxylation activity in *CpFAH12*. This site corresponds to I152 in castor, I148 in *O. violaceus*, N149 in *Physaria* and *Paysonia*, and N148 in Arabidopis *FAD2*, while in *H. benghalensis*, this residue was threonine (T148) which was same as all other *FAD2s* (Figure 5). Therefore, the hydroxylases are somewhat more variable in their sequence than the desaturases, with most of the changes not shared by all hydroxylases, suggesting that they are not essential to the hydroxylase function, or that multiple coordinated changes are needed to change from desaturase to hydroxylase.

It is not surprising that the sequence similarity between *Physaria* and *Paysonia* orthologs of the *FAH12* is high, and this is confirmed by the grouping together of the hydroxylase sequences of *Physaria* and *Paysonia* in the phylogeny when compared with the closely related desaturase sequences. This suggests that the hydroxylase gene evolved in the common ancestor of these two genera and is of relatively recent origin in the family. Comparison of the hydroxylase genes and their related *FAD2* desaturase genes in *Paysonia* and *Physaria* indicates that the three recognized and the fourth putative histidine boxes are almost totally conserved, and that there is only one residue, just before the first histidine box, that uniquely separates desaturase and hydroxylase sequences. This change, from alanine to glycine (105G), is chemically conservative, as both amino acids are non-polar with aliphatic residues. The most variable part of the alignment occurs just before or after the last histidine box (296V, 322S, 324I), and is caused by several changes in three amino acids in the hydroxylase sequences. None of the changes, which occur in more than one species, appear to be lineage specific. This is supported by the PAML analyses, which found no evidence of positive selection for the branch leading to the *Paysonia* and *Physaria* hydroxylases, but rather evidence of relaxed selection compared to the rest of the desaturase tree, consistent with changes in only a few amino acids. The molecular phylogeny confirms the close evolutionary relationship between desaturases and hydroxylases, making it likely that it will be possible to alter gene sequence and thus change the desaturase activity of desaturases from other Brassicaceae to hydroxylases. However, as previous studies have shown (Smith et al., 2000;Smith et al., 2003), just changing the activity of the desaturase is insufficient for significant accumulation in TAG, and that changing the activity of a coordinated set of genes will be necessary to materially change oil content in HFA-free Brassicaceae species. We hope that the transcriptomes we have produced from the sister genus to *Physaria* will help in elucidating what gene modifications will be needed to effectively engineer new sources of HFAs for the HFA oil industry.

## 5 Conclusions

In this study, we applied RNA-Seq to investigate the seed developmental transcriptome of *Paysonia*. A total of 245 million cleaned pair-end reads were obtained and used for the de novo transcriptome assembly with the Trinity program, resulting in a total of 122,730 contigs with an average length of 1,255 bp, including the transcripts of all major enzymes involved in FA biosynthesis and metabolism. A close look at the dynamics of FA/HFA biosynthesis-related genes revealed a similarity of HFA synthesis between *Paysonia* and *Physaria*. Analyses of hydroxylases and related desaturases show their close relationship but do not clearly show conserved changes that convert desaturates to hydroxylases. We believe that this data will provide a valuable resource for future investigations of the HFA synthesis in *Paysonia* and *Physaria*.

## 7 Author Contributions

AND designed the research and secured funding. AS and MM-H performed morphological analyses. GB and JB tested markers and performed phylogenetic analyses. HH performed bioinformatics and gene expression analyses. HH and AND interpreted the data and wrote the manuscript.

## 9 Acknowledgments

We wish to thank members of the Doust lab for plant care and crossing of *Paysonia* accessions. Lipid analysis and advice was provided by Russell Williams at the Proteomics and Mass Spectrometry Facility at the Donald Danforth Plant Science Center. Sequencing support was provided by the Genomics Core Facility at West Virginia University, the BioHPC Lab at Cornell University, and the Oklahoma State University Genomics and Proteomics Center.

