## Supplementary Material for "Transcriptomic analysis of seed development in *Paysonia auriculata* (Brassicaceae) identifies genes involved in hydroxy fatty acid biosynthesis and seed maturation"

### 1 Supplementary Figures and Tables

#### 1.1 Supplementary Figures

**Supplementary Figure 1.** Seed area of individual siliques collected at different times over development. Siliques were collected from weeks 2 to 8 after pollination. Each mean and confidence interval represent a single silique. Error bars represent 95% confidence intervals.

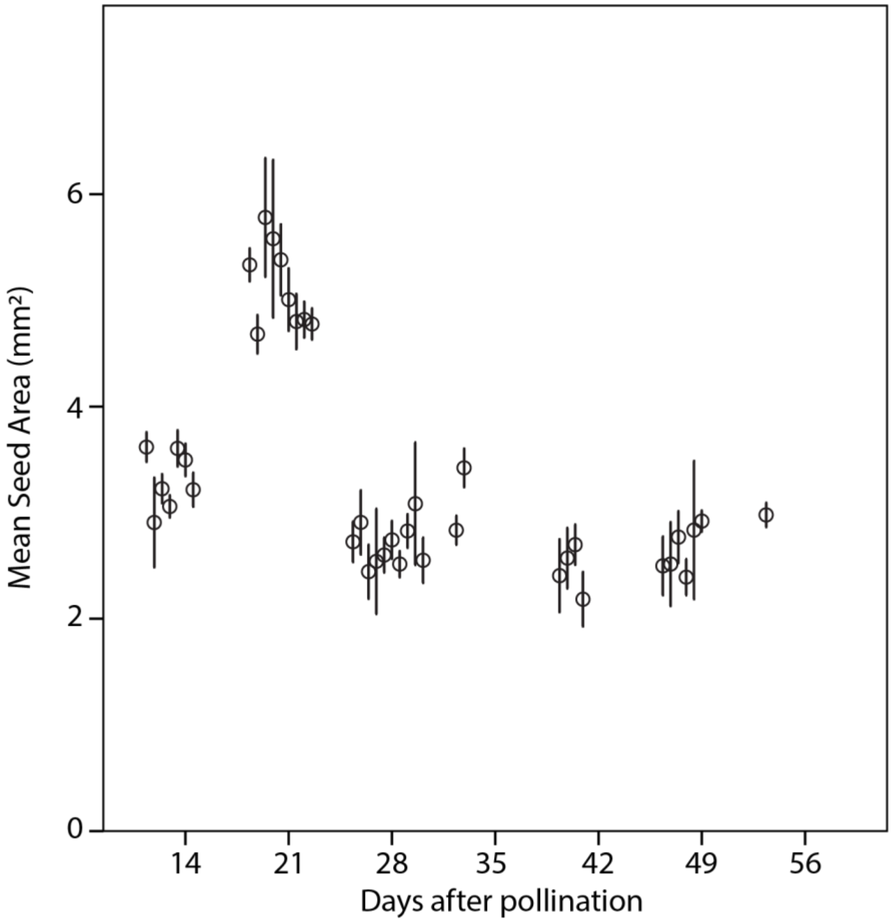

**Supplementary Figure 2.** Seed number of individual siliques collected at different times over development. Filled in circles are actual numbers in individual siliques, open diamond is mean, bars and whiskers are 95% confidence intervals.

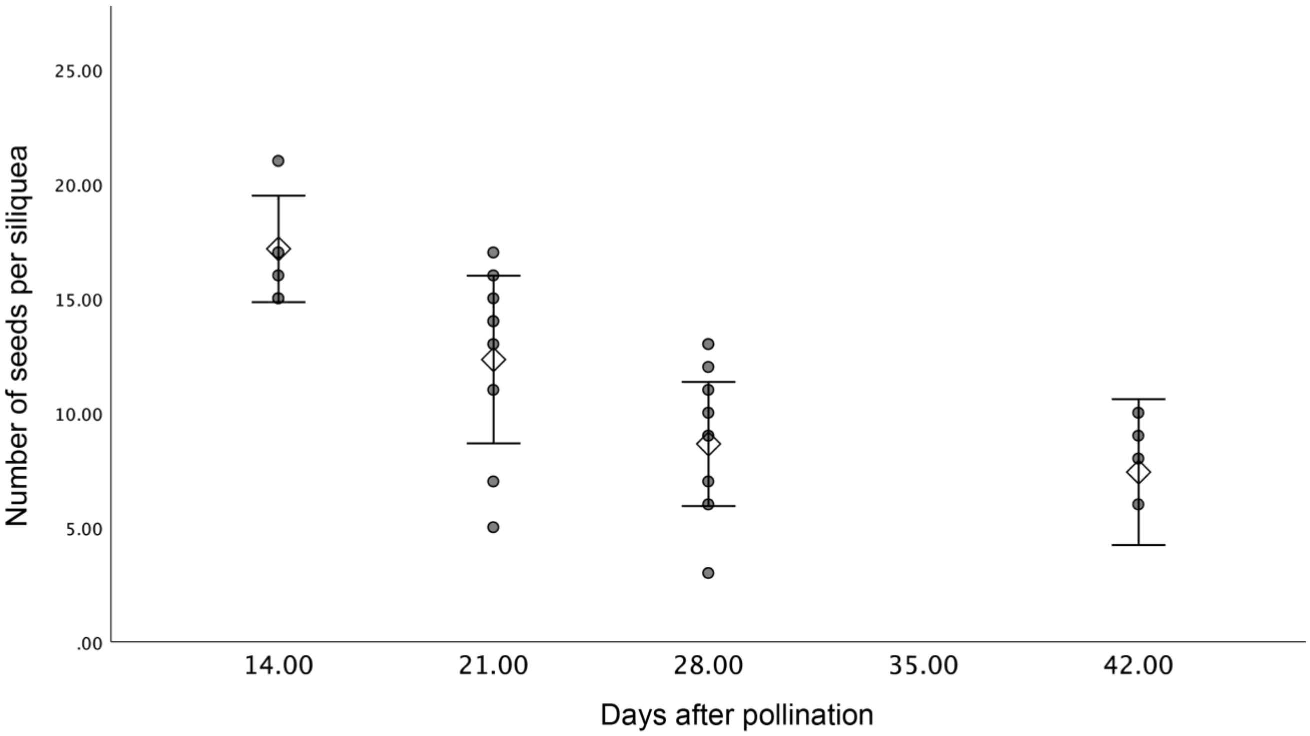

**Supplementary Figure 3.** Phylogenetic tree of FAH12 in various species.

PfFAH12 (AAC32755), PlFAH12 (ABQ01458), OvFAD2-2 (APQ40636), HbFAH12-1 (AGN95845), HbFAH12-2 (AGN95846), RcFAH12 (XP_002528127), CpFAH12 (BAW27658),

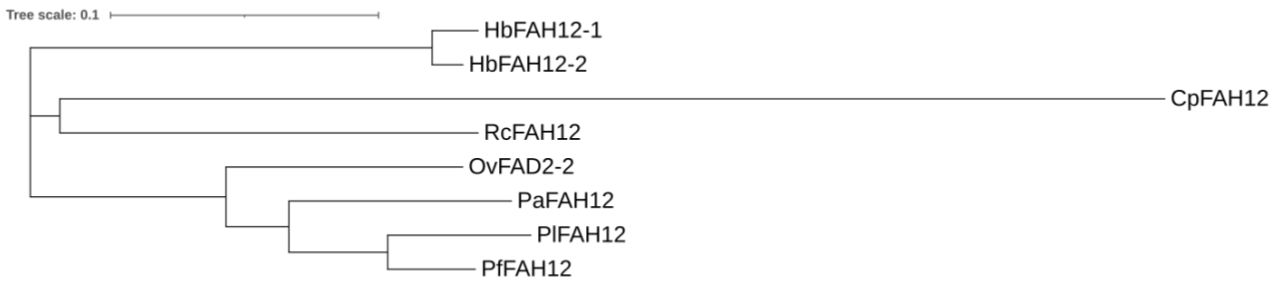

**Supplementary Figure 4.** Phylogenetic tree of FAD2 in various species.

PfFAD2 (AFP21683), AtFAD2 (NP_001319529), OvFAD2-1 (APQ40635), HbFAD2-1 (AGN95841), HbFAD2-2 (AGN95842), HbFAD2-3 (AGN95843), RcFAD2 (NP_001310648), CpFAD2 (ABS18716).

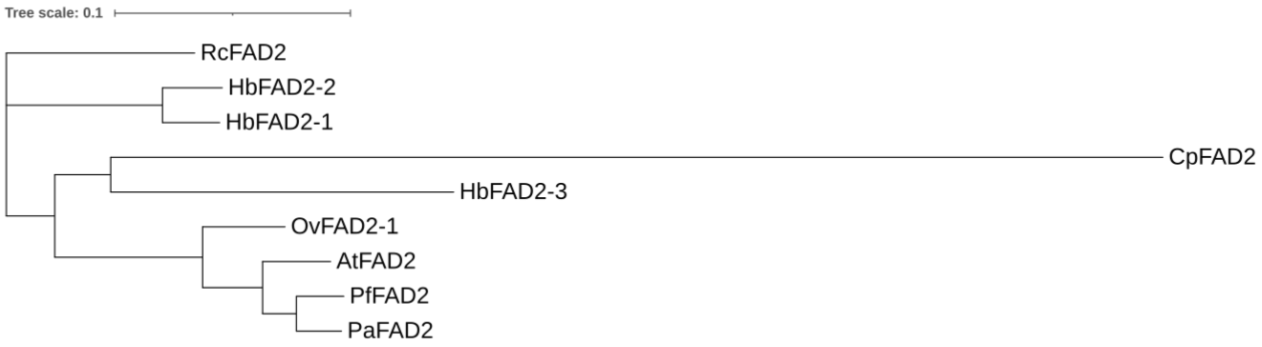

**Supplementary Figure 5.** Phylogenetic tree of FAD3 in various species.

PfFAD3-1 (AWX65624), PfFAD3-2 (AWX65625), PlFAD3-1 (UIH10881), PlFAD3-2 (UIH10882), AtFAD3 (AT2G29980), CsFAD3 (ABA55806), OvFAD3 (AWY93817), RcFAD3 (EEF36775).

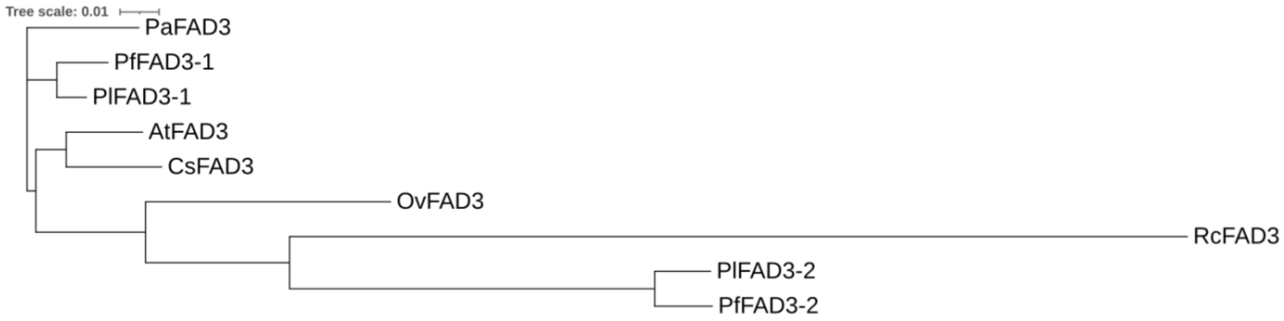

**Supplementary Figure 6.** Amino acid alignment of FAD3 in various species.

PfFAD3-1 (AWX65624), PfFAD3-2 (AWX65625), PlFAD3-1 (UIH10881), PlFAD3-2 (UIH10882), AtFAD3 (AT2G29980), CsFAD3 (ABA55806), OvFAD3 (AWY93817), RcFAD3 (EEF36775). His-boxes were highlighted in yellow.

```
                1                                                                                                  100
PaFAD3          MVVAM-----------------------------------DKRS----------------------------NVNGDS-GAGDPKKG-------------
PlFAD3-1        MVVAM-----------------------------------DQRS----------------------------NVNGDL-SAGDPKKE-------------
PlFAD3-2        MAVTV-----------------------------------DQRS----------------------------DVKGDS------KKE-------------
PfFAD3-1        MVVAM-----------------------------------DKRS----------------------------NVKGDPGGAGDPKKE-------------
PfFAD3-2        MAVTV-----------------------------------DQRS----------------------------DV----------KKE-------------
CsFAD3          MVVAM-----------------------------------DKRS----------------------------NVNGDP-GAGDQKKK------------G
AtFAD3          MVVAM-----------------------------------DQRT----------------------------NVNGDP-GAGDRKKE-------------
OvFAD3          MVVAV-----------------------------------DQRS----------------------------NVNGDS-GASA-RKE-------------
RcFAD3          MAAGWVLSECGLRPLPRIYSRPRIGFTSKTTNLLKLRELPDSKSYNLCSSFKVSSWSNSKQSNWALNVAVPVNVSTVS-GEDDREREEFNGIVNVDEGKG

PaFAD3          EGFDPSAQPPFKIGDIRAAIPKHCWIKSPLRSMSYVVRDICAVAFLAIAAVYFDSWFLWPLYWAAQGTLFWAIFVLGHDCGHGSFSDIPLLNSLVGHILH
PlFAD3-1        ERFDPSAQPPFKIGDIRAAIPKHCWVKSPLRSMSYVVRDIIAVAFLAIAAVYFDSWFLWPLYWVAQGTLFWAIFVLGHDCGHGSFSDIPLLNSVVGHILH
PlFAD3-2        EAFDPSTKPPFKLGDIRAAIPKQCWVKSPLRSMSYVFRDFSAIAALAFAAVHFDSWFLWPVYWVAQGTLFWALHVLGHDCGHGSFSDIPLLNSVVGHILH
PfFAD3-1        ERFDPSAQPPFKIGDIRAAIPKHCWVKSPLKSMSYVVRDILAVAFLAIAAVYFDSWILWPLYWVAQGTLFWAIFVLGHDCGHGSFSDIPLLNSVVGHILH
PfFAD3-2        EAFDPSAPPPFKLGDIRAAIPKQCWVKSPLRSLSYVFRDFSAIAALAFAAVHFDSWFLWPVYWVAQGTLFWALHVLGHDCGHGSFSDSPLLNSVVGHILH
CsFAD3          EGFDPSAQPPFKIGDIRAAIPKHCWVESPLRSMSYVVRDICAVAALAVAAVYINSWFLWPLYWAAQGTLFWAIFVLGHDCGHGSFSDIPLLNSVVGHILH
AtFAD3          ERFDPSAQPPFKIGDIRAAIPKHCWVKSPLRSMSYVVRDIIAVAALAIAAVYVDSWFLWPLYWAAQGTLFWAIFVLGHDCGHGSFSDIPLLNSVVGHILH
OvFAD3          EGFDPSAQPPFKIGDIRAAIPKYCWVRSPLRSMSYVARDICAVAALAIAALYFDSWIIWPLYWAAQGTLFWAIFVLGHDCGHGSFSDIPLLNSVVGHIIH
RcFAD3          EFFDAGAPPPFTLADIRAAIPKHCWVKNPWRSMSYVLRDVVVVFGLAAVAAYFNNWVAWPLYWFCQGTMFWALFVLGHDCGHGSFSNNPKLNSVVGHLLH

PaFAD3          SFILVPYHGWRISHRTHHQNHGHVEHDESWVPLPEKVYKNLPHSTRMLRYTVPL-PMLAYPLYLFYRSPGKEGSHFNPYSSLFAPSERKLIATSTICWSI
PlFAD3-1        SFILVPYHGWRISHRTHHQNHGHVENDESWVPLPEKVYKNLPHSTRMLRYTVPL-PMLAYPLYLCYRSPGKEGSHFNPYSSLFAPSERKLIATSTTCWSI
PlFAD3-2        SFILVPFHGWRISHKIHHQNHGHVENDESWVPVSEKVYKNMSRTTRMLRYTLPIFSMLVYPLYLWKRSPGKEGSHFNPNSSLFTPKERKLVITSTTCWCL
PfFAD3-1        SFILVPYHGWRISHRTHHQNHGHVENDESWVPLPEKVYKNLPHSTRMLRYTVPL-PMLAYPLYLCYRSPGKEGSHFNPYSSLFAPSERKLIATSTTCWSI
PfFAD3-2        SFILVPFHGWRISHKLHHQNHGHVENDESWVPVSEKVYKNMSRTTRMLRYTLPIFSMLVYPLYLWKRSPGKEGSHFNPNSSLFTPKERNLVITSTTCWCL
CsFAD3          SFILVPYHGWRISHRTHHQNHGHVENDESWVPLPERVYKNLPHSTRMLRYTVPL-PMLAYPLYLCYRSPGKEGSHFNPYSSLFAPSERKLIATSTTCWSI
AtFAD3          SFILVPYHGWRISHRTHHQNHGHVENDESWVPLPERVYKKLPHSTRMLRYTVPL-PMLAYPLYLCYRSPGKEGSHFNPYSSLFAPSERKLIATSTTCWSI
OvFAD3          SFILVPYHGWRISHRTHHQNHGHVENDESWVPLPKKLYKNLPYNERMFRYTVPL-PMLAYPLYLWYRCPGKKGSHFNPYSSLFDPSERKLIATSTTCWSI
RcFAD3          SSILVPYHGWRISHRTHHQNHGHVENDESWHPLSEKIFKSLDNVTKTLRFSLPF-PMLAYPFYLWSRSPGKKGSHFHPDSGLFVPKERKDIITSTACWTA

PaFAD3          MLLTLITLSFIFGPISVLKVYGVPYIIFVMWLDAVTYLHHHGHDEKLPWYRGKEWSYLRGGLTTIDRDYGIFNNIHHDIGTHVIHHLFPQIPHYHLVDAT
PlFAD3-1        MFVTLIALSFIFGPISVLKVYGVPYIIFVMWLDAVTYLHHHGHDEKLPWYRGKEWSYLRGGLTTIDRDYGIFNNIHHDIGTHVIHHLFPQIPHYHLVEAT
PlFAD3-2        MLATLIALSYIFGPLLVLKVYGVPYIISVMWLDAVSYLHHHDHDEKLPWYRGKEWSYLRGGLTTIDRDYGIFNEVHHYCGTHVIHHLFPQIPHYHLVEAT
PfFAD3-1        MFVTLIALSFIFGPISVLKVYGVPYIIFVMWLDAVTYLHHHGHDEKLPWYRGKEWSYLRGGLTTIDRDYGIFNNIHHDIGTHVIHHLFPQIPHYHLVEAT
PfFAD3-2        MLATLIALSYIFGPVSVLKVYGVPYIISVMWLDAVSYLHHHDHDEKLPWYRGKEWSYLRGGLTTIDRDYGIFNEVHHYCGTHVIHHLFPQIPHYHLVEAT
CsFAD3          MFVTLIALSFIFGPLSVLKVFGVPYIIFVMWLDAVTYLHHHGHDEKLPWYRGKEWSYLRGGLTTIDRDYGIFNNIHHDIGTHVIHHLFPQIPHYHLVDAT
AtFAD3          MFVSLIALSFVFGPLAVLKVYGVPYIIFVMWLDAVTYLHHHGHDEKLPWYRGKEWSYLRGGLTTIDRDYGIFNNIHHDIGTHVIHHLFPQIPHYHLVDAT
OvFAD3          MVATLIYLSSLVGPVTVLKVYGVPYIIFVMWLDTVTYLHHHGHNEKLPWYRGKEWSYLRGGLTTIDRDYGIFNNIHHDIGTHVIHHLFPQIPLYHLVEAT
RcFAD3          MAALLVYLNFSMGPVQMLKLYGIPYWIFVMWLDFVTYLHHHGHEDKLPWYRGKAWSYLRGGLTTLDRDYGWINNIHHDIGTHVIHHLFPQIPHYHLVEAT

PaFAD3          KAAKHVLGRYYREPKTSGAIPIHLVESLVASIKKDHYVSDSGDIVFYQTDPDLYVVYASDKSKIN
PlFAD3-1        KAAKHVLGRYYREPKTSGAIPIHLVESLVASIKKDHYVSDSGDIVFYETDPDLYVVYASDKSKIN
PlFAD3-2        NAAKHVMGKYYREPKKSGAIPVHLVESLFASLKKNHYVSDTGDIVFYKTDPDLQDYAYDKSKLT
PfFAD3-1        KAAKHVLGRYYREPKTSGAIPIHLVESLVSSIKKDHYVSDSGDIVFYETDPDLYVVYASDKSKIN
PfFAD3-2        NAAKHVMGKYYREPKKSGAIPVHLVESLFASLKKNHYVSDTGDIVFYKTDPDLQDYAYDKTKVN
CsFAD3          KSAKHVLGRYYREPQTSGAIPIHLVESLVASIKKDHYVSDTGDIVFYETDPDLYVVYASDKSKIN
AtFAD3          KAAKHVLGRYYREPKTSGAIPIHLVESLVASIKKDHYVSDTGDIVFYETDPDLYVVYASDKSKIN
OvFAD3          KAAKHVLGRYYREPKTSGAIPIHLVESLVASLKKDHYVSDTGDVVFYETDPDLYGYASEKSKIN
RcFAD3          EAAKPVMGKYYREPKKSGPLPLHLLGSLVRSMKEDHYVSDTGDVVYYQKDPKLSGIGGEKTE--
```

**Supplementary Figure 7.** Phylogenetic tree of KCS18 in various species.

CsKCS18-A (ADN10814), CsKCS18-B (ADN10812), CsKCS18-C (ADN10813), AtKCS18 (AT4G34520), PfKCS18 (AAK62348), RcKSC18 (XP_002516167), OvKCS18 (APQ40637).

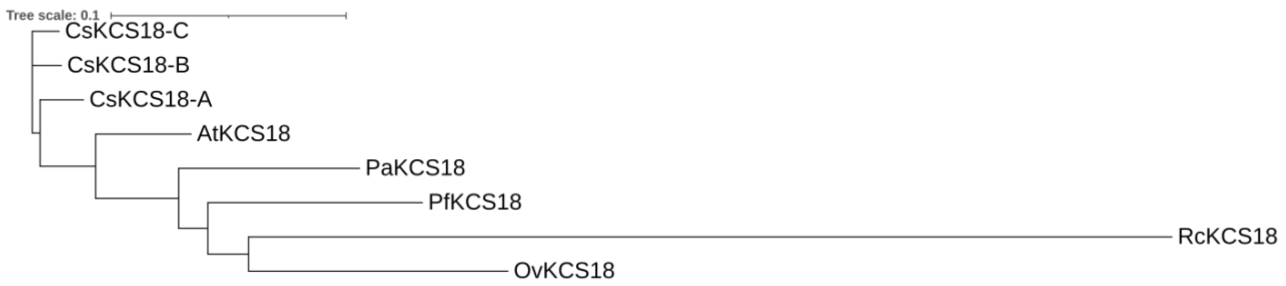

**Supplementary Figure 8.** Amino acid alignment of KCS18 in various species.

CsKCS18-A (ADN10814), CsKCS18-B (ADN10812), CsKCS18-C (ADN10813), AtKCS18 (AT4G34520), PfKCS18 (AAK62348), RcKSC18 (XP_002516167), OvKCS18 (APQ40637).

The colored regions indicated conserved amino acids or sequences, a Gly-Gly region containing a putative active Cys site was highlighted in blue.

```
                    1                                                                                                  100
PaKCS18             MT----------------------------------------SVN---VKLLYHYVLTNFFNLSLFPLTVLI-AGKASQLTENDILLFF-THLQDNLITV
PfKCS18             MT----------------------------------------SLN---IKLLYHYILTNLFNLCLFPLTAFL-AGKASHLTKSDLLMFL-SHLQDNLITV
CsKCS18-A           MT----------------------------------------SVN---AKLLYHYVLTNFFNLCLFPLTALL-AGKASTLTTNDLYHFY-SHLQHNLVTV
CsKCS18-B           MT----------------------------------------SVN---AKLLYHYVLTNFFNLCLFPLTALL-AGKASRLTSNDLYHFY-SHLQHNLITV
CsKCS18-C           MT----------------------------------------SVN---AKLLYHYVLTNFFNLCLFPLTALL-AGKASKLTANDLYHFY-SHLQHNLITV
AtKCS18             MT----------------------------------------SVN---VKLLYRYVLTNFFNLCLFPLTAFL-AGKASRLTINDLHNFL-SYLQHNLITV
OvKCS18             MT----------------------------------------SVN---VKLVYHYIITNFFYLCFFPLTAIV-AGKASRLTLNDLHHFYYSHLQHNLITV
RcKCS18             MANERDLLSTEIVNRGIESSGPNAGSLTFSVRVRRRLPDFLQSVNLKYVKLGYHYLINHAIYLATIPVLVLVFSAEVGSLSREELWRKLWEDARYDLATV

PaKCS18             IVLITFTIFGLVLYIVTRPKSVYLVDYSCYLPPPHLKVSVSSIMDVFYQLRKADP-LRNVACDDPSWFDFLRKIQERSGLGDETYISPGLIQVPPQKTFA
PfKCS18             IVLFTFTIFCLVLYIVTKPKQIYLVDYSCYLPPDHLKVSISSVMDIFYELRKVDP-LCEVGCDDSS-LEFMRKVLERSGLGDETYVPLGLHQVPPQKTFA
CsKCS18-A           ILLFAFSSFGLVLYVVTRRRPVYLVDYSCYLPPPHLKVSVSKVMDIFYQIRKADT-SRNVACDDPSSLDFLRKIQERSGLGDETYSPPGLIHVPPQKTFA
CsKCS18-B           ILLFAFTAFGLVLYIVTRPKPVYLVDYSCYLPPPHLKVSVSKAMDIFYQIRKADT-SRNVACDDPSSLDFLRKIQERSGLGDETYSPQGLINVPPRKTFA
CsKCS18-C           ILLFAFTAFGLVLYIVTRPKPVYLVDYSCYLPPPHLKVSVSKAMDIFYQIRKADT-SRNVACDDPSSLDFLRKIQERSGLGDETYSPQGLINVPPQKTFA
AtKCS18             TLLFAFTVFGLVLYIVTRPNPVYLVDYSCYLPPPHLKVSVSKVMDIFYQIRKADTSSRNVACDDPSSLDFLRKIQERSGLGDETYSPEGLIHVPPRKTFA
OvKCS18             TLLSSFTVFGSLLYIFTRPKPVYLVDYACYLPPPHLKASISRVMDIFYQVRQADP--TAV-CEASS-LEFLRKVQERSGLGDETYGTEGLLQNPPRKNFT
RcKCS18             LSFFGVLVFTVSAYFMSRPRSIYLLDFACYRPHDDLKVTKDQFMEM---LRKSGK------YDEAS-LEFQKRILQSSGIGDETYIPKAVMLQENCTTMK

PaKCS18             AVREETEMVIIGALENLFENTKVNPREIGILVVNSSVFNPTPSLCAMIVNTFKLRSNIKSVNMGGMGCSAGVIAIDLAKDLLHVHKNTYALVVSTENTTI
PfKCS18             AIKDETEQVIKGALENLFENTKVNPREIGILVINSSMFNPTPSLSAMVINTFKLRSNIKSFNLGGMGCSAGVIAIDLAKDLLHVHKNTYALVISTENITI
CsKCS18-A           ASREETEQVIIGALEKLFENTKVNPREIGILVVNSSMFNPTPSLSAMVVNTFKLRSNIKSFSLGGMGCSAGVIAIDLAKDLLHVHKHTYALVVSTENITQ
CsKCS18-B           ASREETEQVIIGALDKLFENTKVNPREIGILVVNSSMFNPTPSLSAMVVNTFKLRSNIKSFSLGGMGCSAGVIAIDLAKDLLHVHKNTYALVVSTENITQ
CsKCS18-C           ASREETEQVIIGALEKLFENTKVNPREIGILVVNSSMFNPTPSLSAMVVNTFKLRSNIKSFSLGGMGCSAGVIAIDLAKDLLHVHKNTYALVVSTENITQ
AtKCS18             ASREETEKVIIGALENLFENTKVNPREIGILVVNSSMFNPTPSLSAMVVNTFKLRSNIKSFNLGGMGCSAGVIAIDLAKDLLHVHKNTYALVVSTENITQ
OvKCS18             AAREETEQVITGALEKLFKNTKVNTKEIGILVVNSGMFNPTPSLSAMVVNKFKLRSNIRSFNLGGMGCSASVIAVDLAKDLLQVHKNTYALVVSTENITY
RcKCS18             EGRLEASSVIFGALDELFEKTRVRPKDVGVLVVNCSIFNPTPSLSAMVINHYKMRGNILSYNLGGMGCSAGIIAVDLARDMLQANPNNIAVVVSTEMVGY

PaKCS18             NAYAGENRSMMMGNCLFRVGGAAILLSNKPGDRRRSKYKLVHTVRTNTGADDNSFRCVQQQEDEMGKLGVSLSKDLINVAGTAVKKNISTLGPLILPLSE
PfKCS18             TAYAGENRSMNVSNCLFRIGGAAILLSNKPRDKRRSKYKLAHTVRTHTGADDMSYRCVQQEEDEMGKVGVRLSKDITTVAGTAVKKNISTLGPLILPLSE
CsKCS18-A           GIYAGENRSMMVSNCLFRVGGAAILLSNKPGDRRRSKYKLCHTVRTHTGADDKSFRCVQQGDDESGKIGVCLSKDITVVAGTALKKNIATLGPLILPLSE
CsKCS18-B           GIYAGENRSMMVSNCLFRVGGAAILLSNKPGDRRRSKYKLCHTVRTHTGADDMSFRCVQQGDDESGKIGVCLSKDITVVAGIALKKNIATLGPLILPLSE
CsKCS18-C           GIYAGENRSMMVSNCLFRVGGAAILLSNKLGDRRRSKYKLCHTVRTHTGADDKSFRCVQQGDDEGGKIGVCLSKDITVVAGTALKKNIATLGPLILPLSE
AtKCS18             GIYAGENRSMMVSNCLFRVGGAAILLSNKSGDRRRSKYKLVHTVRTHTGADDKSFRCVQQEDDESGKIGVCLSKDITNVAGTTLTKNIATLGPLILPLSE
OvKCS18             NSYTGDNKSMMVSNCLFRVGGAAILLSNKPRERRRSKYELVHTVRTHAGADDMSYRCVIQEDDESGKNGVSLSKDITTVAGRAITINMSTLGPLVLPISE
RcKCS18             NWYPGKERSMMIPNCFFRMGCSAVLLSNRRRDYRRAKYRLEHIVRTHKGADDRSFRSVYQDEDEQKFKGLKVSKELMEIGGDALKTNITTLGPLVLPFSE

PaKCS18             KFLFFVNFIAKKLLKDKIKHN-------YVPDFKLAIDHFCIHAGGRGVLDVLEKNLRLSTIDVEASRSTLHRFGNTSSSSIWYELAYIEAKGRMKKGNR
PfKCS18             KLLYFVSFIAKKLLKEKIKNY-------YVPDLKLAINHFCIHAGGRGVLDVLEKNLRLSPIDVEASRSTLHRFGNTSSSSIWYELAYIEAKGRMKKGDK
CsKCS18-A           KFLFLVTFIAKKLLKDKIKHY-------YVPDFKLAIDHFCIHAGGRAVIDVLEKSLGLSPIDVEASRSTLHRFGNTSSSSIWYELAYIEAKGRMKKGNR
CsKCS18-B           KFLFLVTFIAKKLLKDKIKHY-------YVPDFKLAIDHFCIHAGGRAVIDVLEKSLGLSPIDVEASRSTLHRFGNTSSSSIWYELAYIEAKGRMKKGNR
CsKCS18-C           KFLFLVTFIAKKLLKDKIKHC-------YVPDFKLAIDHFCIHAGGRAVIDVLEKSLGLSPIDVEASRSTLHRFGNTSSSSIWYELAYIEAKGRMKKGNR
AtKCS18             KFLFFATFVAKKLLKDKIKHY-------YVPDFKLAVDHFCIHAGGRAVIDELEKNLGLSPIDVEASRSTLHRFGNTSSSSIWYELAYIEAKGRMKKGNK
OvKCS18             KILFFLSLIGKKLMKDKIKNL-------YVPDFKLAIDHFCIHAGGRLVIDVLEKNLGLAPIDVEASRSTLHRFGNTSSSSIWYELAYIEAKGRMKKGNK
RcKCS18             QLIFFVSLVYRQLFGNKKGSLSSPSAKPYIPDYKLAFEHFCVHGASKVVLDEIQRNLELSEKNMEASRMALHRFGNTSSSSIWYELAYLEAKERVKRGDR

PaKCS18             AWQIALGSGFKCNSAVWVALRNVKASANSPWEHCIDRYPVS----ELTKSDTCVQNGRS
PfKCS18             AWQIALGSGFKCNSAVWVALRNVKASANSPWEHCIDRYPV---------PDTCVENGKS
CsKCS18-A           AWQIALGSGFKCNSAVWVALCNVKASANSPWEHCIDRYPVQIDS-DSSKSETHVKNGRS
CsKCS18-B           -GSSKSDTHVKNGRS
CsKCS18-C           AWQIALGSGFKCNSAVWVALCNVKASANSPWEDCIDRYPVQIDS-DSSKSETHVKNGRS
AtKCS18             AWQIALGSGFKCNSAVWVALRNVKASANSPWQHCIDRYPVKIDS-DLSKSKTHVQNGRS
OvKCS18             VWQIALGSGFKCNSAVWVALRNVKASTNSPWEHCIDRYPVNIDSDDSAKPETRVQNGSS
RcKCS18             VWQIAYGSGFKCNSLVWKAIRRVKKPSRNPWLHCIDRYPVQL-----------------
```

**Supplementary Figure 9.** Phylogeny of FAD2 and FAH12 genes in the family Brassicaceae.

FAH12 sequences are in bold, and the branch leading to the FAH12 clade is in red. Length of the tree is indicated by substitution rate at base. Support values are from 1000 bootstrapped analyses. Sequences used in this analysis: FAD2: *A. helleri* (Araha.1578s0006.1), *A. thaliana* (NM_112047.4), *B. rapa* (Brara.E02874.1), *B. stricta* (Bostr.19424s1149.1), *C. glabrata* (KJ573490.1), *C. grandiflora* (Cagra.0430s0042.1), *C. rubella* (Carubv10013933m), *C. sativa* (GU929417.1), *C. sativus* (Cucsa.242720.2), *C. sinensis* (orange1.1g016781m), *E. salsugineum* (Thhalv10020914m), *F. vesca* (mrna22642.1-v1.0-hybrid), *L. campestre* (FJ907546.1), *P. auriculata* (MT496776), *P. fendleri* (KC972621.1), *S. alba* (KY305535.1), *T. hassleriana* (XM_010559645.1). FAH12: *P. auriculata* (MT496774), *P. fendleri* (KC972619.1), *P. lindheimeri* (EF432246.1).

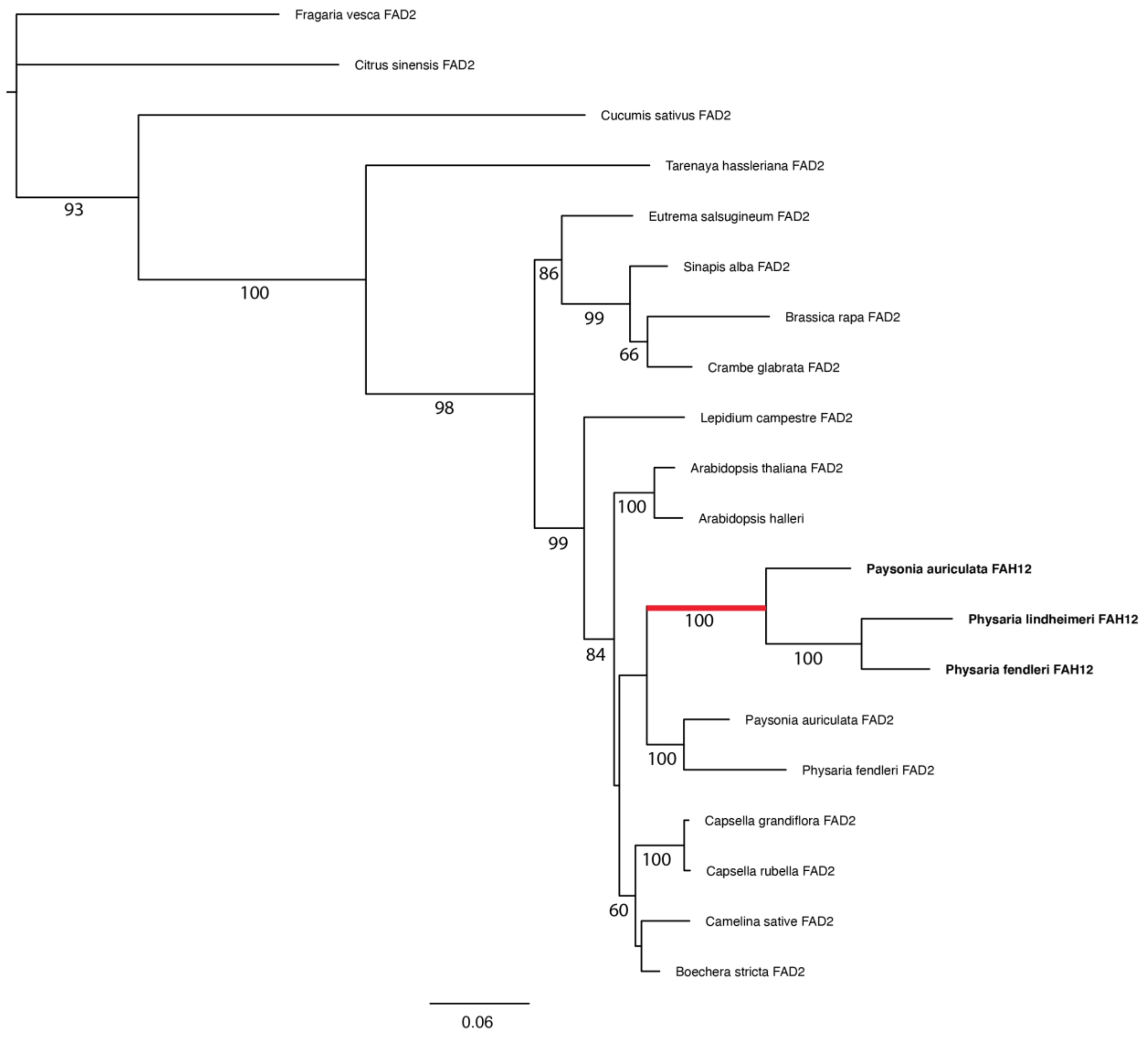

### 1.2 Supplementary Tables

**Supplementary Table 1. Summary statistics of sequencing data of individual library**

| Sample | Number of raw reads | Number of filtered reads | Overall alignment rate to Trinity assembly using Bowtie2 |
|---|---|---|---|
| Pa_2_rep1 | 39,905,282 | 21,876,307 | 82.38% |
| Pa_2_rep2 | 31,471,350 | 16,989,546 | 82.43% |
| Pa_2_rep3 | 38,260,148 | 22,229,570 | 81.02% |
| Pa_3_rep1 | 37,529,310 | 22,195,302 | 78.40% |
| Pa_3_rep2 | 41,054,622 | 23,754,920 | 79.20% |
| Pa_3_rep3 | 39,799,045 | 21,283,284 | 77.88% |
| Pa_4_rep1 | 38,615,229 | 20,826,262 | 78.78% |
| Pa_4_rep2 | 37,044,630 | 20,595,350 | 80.17% |
| Pa_4_rep3 | 33,715,960 | 17,721,020 | 79.40% |
| Pa_6_rep1 | 34,663,919 | 19,231,331 | 78.35% |
| Pa_6_rep2 | 39,465,750 | 19,866,715 | 83.25% |
| Pa_6_rep3 | 32,607,593 | 18,749,052 | 78.08% |

**Supplementary Table 2. Summary statistics of *de novo* transcriptome assembly with Trinity**

| Trinity *de novo* assembly | |
|---|---|
| Total assembled bases | 226,807,152 |
| Number of transcripts | 270,114 |
| Number of isoforms | 122,730 |
| Average transcript length | 840 bp |
| Max gene length | 18,345 bp |
| Min gene length | 201 bp |
| Transcript N50 | 1,255 bp |
| GC content | 40.80 |
| Length and number distribution of isoforms | |
| 200 – 400 | 67,710 |
| 400 – 600 | 17,198 |
| 600 – 800 | 9,376 |
| 800 – 1,000 | 6,408 |
| 1,000 – 1,500 | 10,313 |
| > 1500 | 11,725 |
| Total | 122,730 |

**Supplementary Table 3. GO enrichments of DE genes from different clusters in SOM.**

| GO | Term | Cluster 1 | Cluster 2 | Cluster 3 | Cluster 4 |
|---|---|---|---|---|---|
| GO:0009414 | response to water deprivation | 6.4 | 6.5 | 1.5 | 6.9 |
| GO:0009416 | response to light stimulus | 4.3 | 6.0 | 2.4 | 0.4 |
| GO:0055114 | oxidation-reduction process | 14.4 | 19.1 | 8.9 | 6.8 |
| GO:1901700 | response to oxygen-containing compound | 5.2 | 9.4 | 9.2 | 4.1 |
| GO:0000302 | response to reactive oxygen species | 0.5 | 7.7 | 4.1 | 0.3 |
| GO:0006979 | response to oxidative stress | 3.3 | 11.6 | 6.3 | 1.3 |
| GO:0042446 | hormone biosynthetic process | 0.3 | 5.3 | 0.3 | 0.2 |
| GO:0030258 | lipid modification | 0.0 | 14.5 | 0.0 | 0.3 |
| GO:0044242 | cellular lipid catabolic process | 0.0 | 23.8 | 0.3 | 0.1 |
| GO:0006996 | organelle organization | 0.0 | 3.2 | 0.0 | 0.0 |
| GO:0019222 | regulation of metabolic process | 0.0 | 11.2 | 0.0 | 0.0 |
| GO:0080090 | regulation of primary metabolic process | 0.0 | 8.0 | 0.0 | 0.0 |
| GO:0046907 | intracellular transport | 0.0 | 3.8 | 0.0 | 0.0 |
| GO:0015031 | protein transport | 0.0 | 5.0 | 0.0 | 0.0 |
| GO:0007031 | peroxisome organization | 0.0 | 12.2 | 0.0 | 0.0 |
| GO:0007568 | aging | 0.0 | 1.1 | 2.0 | 5.1 |
| GO:0005985 | sucrose metabolic process | 1.0 | 0.1 | 2.2 | 9.3 |
| GO:0055085 | transmembrane transport | 1.6 | 0.4 | 5.9 | 4.1 |
| GO:0010876 | lipid localization | 1.6 | 0.0 | 5.4 | 5.0 |
| GO:0019915 | lipid storage | 0.4 | 0.1 | 5.8 | 2.0 |
| GO:0090351 | seedling development | 0.1 | 0.5 | 7.7 | 2.2 |
| GO:0009737 | response to abscisic acid | 2.3 | 2.2 | 3.6 | 2.0 |
| GO:1901568 | fatty acid derivative metabolic process | 0.2 | 0.2 | 4.9 | 0.0 |
| GO:0010345 | suberin biosynthetic process | 0.0 | 0.1 | 9.1 | 0.0 |
| GO:0033993 | response to lipid | 2.0 | 1.2 | 3.2 | 1.2 |
| GO:0009698 | phenylpropanoid metabolic process | 5.0 | 0.0 | 21.1 | 0.6 |
| GO:0009725 | response to hormone | 6.8 | 1.0 | 7.4 | 1.5 |
| GO:0006629 | lipid metabolic process | 9.7 | 4.8 | 9.0 | 3.3 |
| GO:0008202 | steroid metabolic process | 11.0 | 0.3 | 3.4 | 1.5 |
| GO:1901658 | glycosyl compound catabolic process | 7.7 | 0.0 | 2.2 | 0.0 |
| GO:0071555 | cell wall organization | 14.0 | 0.0 | 3.8 | 0.1 |
| GO:0006633 | fatty acid biosynthetic process | 14.9 | 0.0 | 8.2 | 1.8 |
| GO:0009813 | flavonoid biosynthetic process | 9.3 | 0.2 | 4.4 | 1.3 |
| GO:0008610 | lipid biosynthetic process | 14.6 | 0.2 | 6.3 | 2.8 |
| GO:0006637 | acyl-CoA metabolic process | 5.0 | 0.0 | 1.1 | 1.0 |
| GO:0044264 | cellular polysaccharide metabolic process | 12.6 | 0.0 | 1.2 | 0.9 |
| GO:0006733 | oxidoreduction coenzyme metabolic process | 13.5 | 0.6 | 0.0 | 0.0 |
| GO:0016051 | carbohydrate biosynthetic process | 17.5 | 0.1 | 0.4 | 0.5 |
| GO:0016049 | cell growth | 2.5 | 0.0 | 0.1 | 0.0 |
| GO:0010191 | mucilage metabolic process | 12.1 | 0.0 | 0.1 | 0.0 |
| GO:0010154 | fruit development | 4.2 | 1.5 | 1.7 | 1.4 |
| GO:0016053 | organic acid biosynthetic process | 14.6 | 0.9 | 2.9 | 0.2 |
| GO:0010214 | seed coat development | 9.5 | 0.1 | 1.0 | 0.0 |
| GO:0005976 | polysaccharide metabolic process | 14.9 | 0.0 | 1.8 | 0.3 |
| GO:0019684 | photosynthesis, light reaction | 22.4 | 0.0 | 1.9 | 0.3 |

White to red color keys indicate negative log10-transformed *p*-values for enrichment ($p < 0.05$).

**Supplementary Table 4. Primers used in cloning and qRT-PCR**

| Gene | Sequence (5' – 3') | Purpose |
|---|---|---|
| PaFAH12 | F: CACCATGGGTGCTGGTGGAAGAAT<br>R: TCATAACTTATTGTTGAACCAGTAG | Cloning |
| PaFAD2 | F: CACCATGGGTGCTGGTGGAAGAAT<br>R: TCATAACTTATTGTTGTACCAGTACA | Cloning |
| PaFAD3 | F: CACCATGGTGGTTGCTATGG<br>R: TTAATTGATTTTAGATTTGTC | Cloning |
| PaKCS18 | F: CACCATGACGTCCGTGAACGTA<br>R: TCAAGACCGCCCATTTTGG | Cloning |
| PaFAH12 | F: ATTACTTGGCTTGGCCCCTC<br>R: CGTGACCAATGACCCAGACA | qRT-PCR |
| PaFAD2 | F: CACGCTTTTAGCAACTACCAGT<br>R: TCGAGGGATCCTGTGTTGGA | qRT-PCR |
| PaFAD3 | F: GCGCTGGAGATCCGAAGAAA<br>R: GATCTTAAACGGCGGCTGTG | qRT-PCR |
| PaKCS18 | F: ATCAACAATCGATGTCGAAGCA<br>R: TCCCAATGCAATCTGCCAAATC | qRT-PCR |
| 18S | F: GAGAAACGGCTACCACATCCA<br>R: CCGTGTCAGGATTGGGTAATTT | qRT-PCR |
